# Circuit Analysis of the Drosophila Brain using Connectivity-based Neuronal Classification Reveals Organization of Key Communication Pathways

**DOI:** 10.1101/2022.03.11.484039

**Authors:** Ketan Mehta, Rebecca F. Goldin, Giorgio A. Ascoli

**Affiliations:** Dept. of Bioengineering and Center for Neural Informatics, Structures, and Plasticity, George Mason University, Fairfax, VA 22030, USA; Dept. of Mathematical Sciences and Center for Neural Informatics, Structures, and Plasticity, George Mason University, Fairfax, VA 22030, USA

**Keywords:** fruit fly connectome, neural circuits, dopaminergic hub, cell type classification, functional pathways, multisensory integration

## Abstract

We present a functionally relevant, quantitative characterization of the neural circuitry of *Drosophila melanogaster* at the mesoscopic level of neuron types as classified exclusively based on potential network connectivity. Starting from a large neuron-to-neuron brain-wide connectome of the fruit fly, we use stochastic block modeling and spectral graph clustering to group neurons together into a common “cell class” if they connect to neurons of other classes according to the same probability distributions. We then characterize the connectivity-based cell classes with standard neuronal biomarkers, including neurotransmitters, developmental birthtimes, morphological features, spatial embedding, and functional anatomy. Mutual information indicates that connectivity-based classification reveals aspects of neurons that are not adequately captured by traditional classification schemes. Next, using graph-theoretic and random walk analyses to identify neuron classes as hubs, sources, or destinations, we detect pathways and patterns of directional connectivity that potentially underpin specific functional interactions in the *Drosophila* brain. We uncover a core of highly interconnected dopaminergic cell classes functioning as the backbone communication pathway for multisensory integration. Additional predicted pathways pertain to the facilitation of circadian rhythmic activity, spatial orientation, fight-or-flight response, and olfactory learning. Our analysis provides experimentally testable hypotheses critically deconstructing complex brain function from organized connectomic architecture.

**AUTHOR SUMMARY:** The potential synaptic circuitry of a neural system constitutes the fundamental architectural underpinning of its in vivo dynamics, plasticity, and functions. The fruit fly neural circuit presented here captures the latent stochastic patterns of network connectivity and provides a fundamental parts list for reverse-engineering brain computation. Mapping the interactions among connectivity-based neuronal classes to development, morphology, physiology, and transcriptomics result in testable hypotheses on the relationship between whole-brain neural architecture and behavior.

## INTRODUCTION

While it is well established that the brain is composed of distinct cell types, the extent of this cellular diversity remains a long standing open question in neuroscience (Zeng & Sanes 2017). A popular approach to studying brain computation is to model the nervous system as a giant interconnected network of these distinct cell types, each playing a specific role. The underlying assumption is that the intricate connectivity patterns of the neural network constitute the fundamental architectural underpinning of its in vivo dynamics and functions (Abbott et al. 2020; Jonas & Kording 2015). From this perspective, quantitatively characterizing the distinct cell types and their relation to synaptic circuitry is paramount to deconstructing brain computation.

Recent advancements in data acquisition and imaging techniques have enhanced our ability to construct very large scale maps of the neural circuitry in the form of connectomes. Macroscale connectomes are well suited to map the circuitry across the whole brain by parcellating it into distinct anatomical regions, but lack the ability to trace individual neurons. On the other hand, microscale connectomes enable the mapping of neural circuitry at the level of single neurons, capturing information pertaining to cell bodies, neurites, and individual synapses. These fine details are critical to determine the relation between structure and signal processing in the brain. However, the inherently massive scale of microscopic connectomes is not ideal for directly inferring brain-wide mechanisms and properties. A common practice is to group neurons in a microscale connectome into a common cell type by subsets of multifarious properties, including physiology, biochemistry, and morphology, and subsequently analyze the interactions between these groups. The expedient abundance of data has allowed the creation of increasingly unbiased descriptive taxonomies (DeFelipe et al. 2013; Yuste et al. 2020) for the grouping of neurons. However, these experimentally accessible dimensions are only indirect proxies for the mechanistically more relevant features of network connectivity, developmental control, and experience-dependent plasticity (Armañanzas & Ascoli 2015; Shepherd et al. 2019).

To address these challenges, we construct a mesoscopic level circuit of neuron types as classified exclusively based on their patterns of potential synaptic connectivity. We leverage a recently developed mathematical framework (Mehta et al. 2021), which models a connectome as a directed stochastic block model (SBM) graph, to group neurons together into a common “cell class” if they connect to neurons of other classes according to the same probability distributions. Here we build and expand upon that approach applying it to a 19,902-neuron potential connectome of *Drosophila melanogaster*. The nodes of the identified circuit represent the derived connectivity-based neuronal classes, while the directed edges represent the connection probabilities between neurons in those respective classes.

The *Drosophila* brain has a tractable brain size consisting of approximately 100,000 (Scheffer et al. 2020) to 200,000 (Raji & Potter 2021) neurons, making it an excellent model organism for connectomic analysis. Over the last several years, numerous *Drosophila* studies have contributed to an increased understanding of how certain anatomical regions and neuron types support specific function and behavior. However, the exact underlying mechanics of these functions remain largely unknown, especially when involving multiple modalities and cross-region integration.

To interpret the functional interactions captured by our derived circuit, we map the connectivity-based neuronal classes with traditional neuronal biomarkers, including neurotransmitters, developmental birthtimes, morphological features, spatial embedding, and functional anatomy. In conjunction with these mappings, we employ graph theoretical measures and random walks to analyze the potential connectivity patterns in the circuit, subsequently identifying pathways underpinning specific functions in the *Drosophila* brain. The predicted communication pathways pertain to multi-sensory integration, rhythmic circadian activity, spatial orientation, fight-or-flight response, and olfactory learning. These results demonstrate that the derived circuit captures latent patterns of interaction not revealed by traditional neuron type classification alone. Overall, our analysis provides experimentally testable hypotheses critically deconstructing complex brain function from organized connectomic architecture.

## MATERIALS AND METHODS

### Data Source

We begin with the recently released *Drosophila* neuron-to-neuron brain-wide potential connectome that was constructed by Shih et al. (2020) using fluorescence imaging data from the FlyCircuit v1.2 database. The FlyCircuit database hosts 28,508 individual neurons from the *Drosophila* brain, out of which 22,866 are from adult female specimens. To construct the network, Shih et al. (2020) co-registered all female neurons to a common brain atlas (Chiang et al. 2011) and inferred the directional potential connectivity (Rees, Moradi, & Ascoli 2017) between them by quantifying their axonal-dendritic spatial overlaps (Huang et al. 2019). Neurons that did not establish both afferent (axonal overlaps with dendrites of other neurons) and efferent (dendritic overlaps with axons of other neurons) connections were discarded. The final connectome consisted of the remaining 19,902 female neurons that form a strongly connected network wherein every neuron can potentially reach, and be reached by, all others. For each neuron, the FlyCircuit database also provides the associated neurotransmitter, Gal4 driver line, developmental birthtime, and functional community.

In this work, the above described connectome is represented as a directed and weighted adjacency matrix *A*_*c*_, with non-negative, integral entries 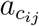 specifying the number of overlapping segment pairs between the axonal arbor of the *i*-th neuron and the dendritic arbor of the *j*-th neuron. Shih et al. (2020) call 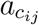 the connection strength. Accordingly, we refer to this connectome as the strength connectome and to *A*_*c*_ as the strength connectivity matrix. *A*_*c*_ is a very sparse matrix with only 0.9% of its entries being non-zero, a maximum connection strength of 1040, and mean connection strength among non-zero entries of 7.8.

### Experimental Design

We stochastically generate multiple binary (unweighted) connectomes from the strength (weighted) connectome. Our functional assumption here is that the structural connectome can be represented as a directed graph with binary edges, i.e., a synaptic connection is either present or absent. Because of the stochastic nature of our model, no two generated binary connectomes are identical and can be considered analogous to connectomes sampled from distinct individual flies. Each generated binary connectome is modeled as a directed (stochastic block model) graph, and clustered using spectral graph clustering. Modeling each graph separately and then merging the individual clustering results via consensus clustering allows for consistent inference. The consensus clusters are used to construct the final circuit. We elaborate on each of these steps below.

#### Stochastic Generation of Binary Connectomes

The strength connectivity matrix is used to derive a connection probability matrix *A*_*p*_, whose entries 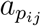 represent the probability that the *i*-th neuron forms a post-synaptic connection with the *j*-th neuron. Intuitively, the larger is the number of overlapping neurites between two neurons, the greater is the chance that they form a synaptic connection. Let *p*_conn_ denote the probability that an overlapping axon-dendritic segment-pair forms a synaptic connection. The connection probability is then derived from the corresponding connection strength using a binomial distribution as follows

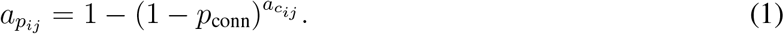

We then proceed to generate multiple binary adjacency matrices *A*^(*ℓ*)^, *l* = 1, 2, …, *G*, each is a stochastic realization of *A*_*p*_. The entry 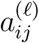 in matrix *A*^(*ℓ*)^ is an independent Bernoulli draw, being either 1 or 0, with corresponding probabilities 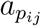and 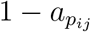, respectively. Further, we trim *A*^(*ℓ*)^ to ensure that it does not have any all-zero rows or columns, i.e., discard any neuron that does not have both a dendritic and axonal connection.

#### Spectral Graph Clustering

Given a *n × n* binary adjacency matrix *A*, we partition the neurons based exclusively on their patterns of stochastic connectivity. In particular, we model the connectome as a SBM graph, and use spectral graph clustering to partition neurons into a common *class* if they connect to neurons in other classes according to the same probability distribution. We refer to these classes as connectivity-based classes.

A SBM graph is parameterized by (i) a block membership probability vector *ρ* = (*ρ*_1_, …, *ρ*_*k*_) in the unit simplex Σ *ρ*_*i*_ = 1, which partitions the *n* vertices into *k* disjoint subsets, and (ii) a *k ×k* block connectivity probability matrix *P*, with entries *p*_*ij*_ *∈* [0, 1]. A SBM assumes that the probability that vertices (neurons) in the *i*-th class form an edge (synaptic connection) with vertices in the *j*-th class can be modeled as an independent Bernoulli trial with parameter *p*_*ij*_.

Recent attempts of applying the SBM framework to model and identify network community structures within small connectomic datasets originating from a variety of sources have yielded considerable success (Betzel, Medaglia, & Bassett 2018; Faskowitz, Yan, Zuo, & Sporns 2018; Jonas & Kording 2015; Moyer et al. 2015; Pavlovic, Vértes, Bullmore, Schafer, & Nichols 2014; Priebe et al. 2017, 2019). Specifically, in Mehta et al. (2021) we developed a mathematical framework that uses SBMs in conjunction with spectral graph clustering to accurately identify connectivity-based classes in large (*≈* 2^12^ *−*2^15^ neurons), and sparse (*≈* 4% edge connectivity) biologically inspired connectomes. Given an artificial surrogate connectome generated using a SBM, the spectral graph clustering was shown to be effective in recovering the true blockmodel structure and accurately assigning each neuron to its respective class, even in the presence of artificially simulated noise (tolerant to as much as 40% pre- and post-synaptic edge misspecification). The clustering framework also scaled extremely well as the number of neurons in the network increases, while being robust over a wide variation in the blockmodel parameters. We now leverage this spectral graph clustering framework to predict the number of connectivity-based classes, and assign each neuron to a class, for an experimentally derived input binary matrix *A*.

The spectral graph clustering is a two-step process comprising of adjacency spectral embedding (ASE) in conjunction with Gaussian mixture model (GMM)-based modeling (Mehta et al. 2021). In the first step, the adjacency matrix is embedded into a lower dimensional space via spectral decomposition of the matrix into its latent vectors. In the second step, the latent vectors are modeled as a Gaussian mixture model (GMM) and clustered using Expectation-Maximization (EM) algorithm.

For any *d ≤* rank(*A*), one can approximate *A* by a rank *d* singular value decomposition

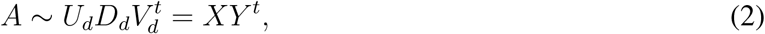

where 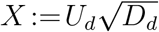 and 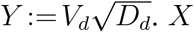 and *Y* are *n × d* matrices, and *D*_*d*_ is a *d × d* diagonal matrix with nonnegative entries called the singular values. For each *A* and choice of embedding dimension *d*, the vectors forming the columns in the augmented matrix **X** :=[*X*|*Y*]^*t*^ provide a dot product embedding of *A* in a 2*d*-dimensional space. The columns of the concatenated matrix **X** are called latent vectors.

The optimal choice of *d* is a known open problem in literature, with no consensus on a best strategy. Choosing a low *d* can result in discarding important information, while choosing a higher *d* than required not only increases computational cost but can adversely effect clustering performance due to the presence of extraneous variables which contribute towards noise in the subsequent statistical inference. We select *d* by using the method outlined in M. Zhu and Ghodsi (2006), which determines the cut-off ‘elbow point’ between relevant and non-relevant dimensions by maximizing a profile likelihood function of the singular values.

For sufficiently dense graphs, and large *n*, the adjacency spectral embedding (ASE) central limit theorem demonstrates that the *n* points in ℝ^2*d*^ behave approximately as random samples from a *k*-component GMM (Athreya et al. 2016). The data is fitted to a GMM using the EM algorithm, which after an initialization of the parameters iteratively improves upon the estimates by maximizing the expected log-likelihood of the probability density function. The accuracy of the EM model fit is known to be sensitive to the initial choices of the parameters. Here, we initialize the EM using parameters obtained by applying model-based hierarchical agglomerative clustering (MBHAC) (Fraley 1998; Scrucca & Raftery 2015) to all *n* data points. Finally, the number of components in the GMM are selected using the Bayesian Information Criterion (BIC), which penalizes the model based on the number of free parameters, i.e., model complexity. The EM model fitting is performed using the mclust R-package (Scrucca, Fop, Murphy, & Raftery 2016).

#### Consensus Clustering using Modified IVC

Let *π*_*ℓ*_ denote the clustering that results from applying the GMM*∘*ASE framework to the corresponding binary adjacency matrix *A*^(*ℓ*)^, for *ℓ* = 1, …, *G*. The clustering *π*_*ℓ*_ maps the *n* vertices *x*_1_, …, *x*_*n*_ into *κ*_*ℓ*_ clusters, such that there is an association of a cluster number *π*_*ℓ*_ (*x*_*i*_) *∈* {1, …, *κ*_*ℓ*_ }, for *i* = 1, …, *n*. In other words, the clustering *π*_*ℓ*_ partitions the vertices into *κ*_*ℓ*_ subsets,

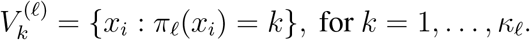

Note that in general each of the clusterings *π*_1_, *π*_2_, …, *π*_*G*_ has different number of clusters *κ*_1_, …, *κ*_*G*_, respectively.

Consensus clustering refers to the process of merging the *G* clusterings into a single, final robust clustering. By combining several clusterings we eliminate any inconsistencies due to noise or outliers, in turn integrating the stochastic variations of individual solutions into a consistent inference. We implement a modified version of the iterative voting consensus (IVC) method introduced by Nguyen and Caruana (2007). The modification removes the reliance on a choice of reference clustering. The two-step process is described below.

**Step 1:** For each clustering *π*_*ℓ*_ we create a corresponding updated version 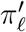 using IVC. Let 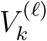 be a cluster of *π*_*ℓ*_. For each *j* = 1, 2, …, *κ*_*ℓ*_, let 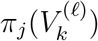 denote the most common label that *π*_*j*_ assigns to the elements of 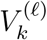 Following Nguyen and Caruana (2007), define the center 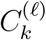of 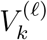 as the sequence of labels associated with 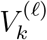 as we vary over the clusterings. While the original method relies on a choice of reference clustering, we iterate sequentially over all *G* clusterings, obtaining a new clustering for each reference clustering. The procedure for obtaining the updated clusterings is detailed in Algorithm 1.

**Step 2:** The final clustering 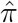 is constructed as follows: we cluster vertices *x* and *x*′ together when they occur in a chain of vertices that are pairwise clustered together at least *τ G* times among the 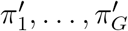 clusterings, where *τ* is a value between 0 and 1, explained below. We then apply a cutoff threshold for a minimal cluster size.

The *similarity matrix* (Nguyen & Caruana 2007) for the clusterings 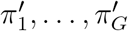 is an *n × n* matrix with the *ij* entry given by the proportion of times that *x*_*i*_ and *x*_*j*_ are clustered together among the clusterings 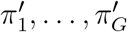. Explicitly, let

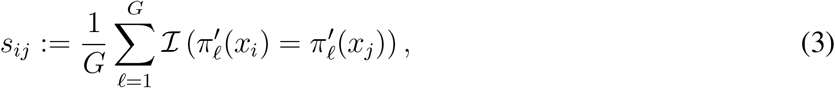

where *ℐ* is the indicator function that is 1 when the statement is true, and 0 otherwise.

We define an equivalence relation that depends on a parameter *τ* among the vertices. We say *x* ∼ *x*′ if there exists a sequence of vertices 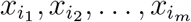 where 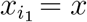 and 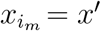, such that 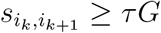 for *k* = 1, …, *m −* 1. One may think of two equivalent vertices as having a chain of vertices between them, each pair along the chain being grouped together at least proportion of *τ* times, among the clusterings 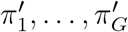.

##### Algorithm 1 IVC over all clusterings

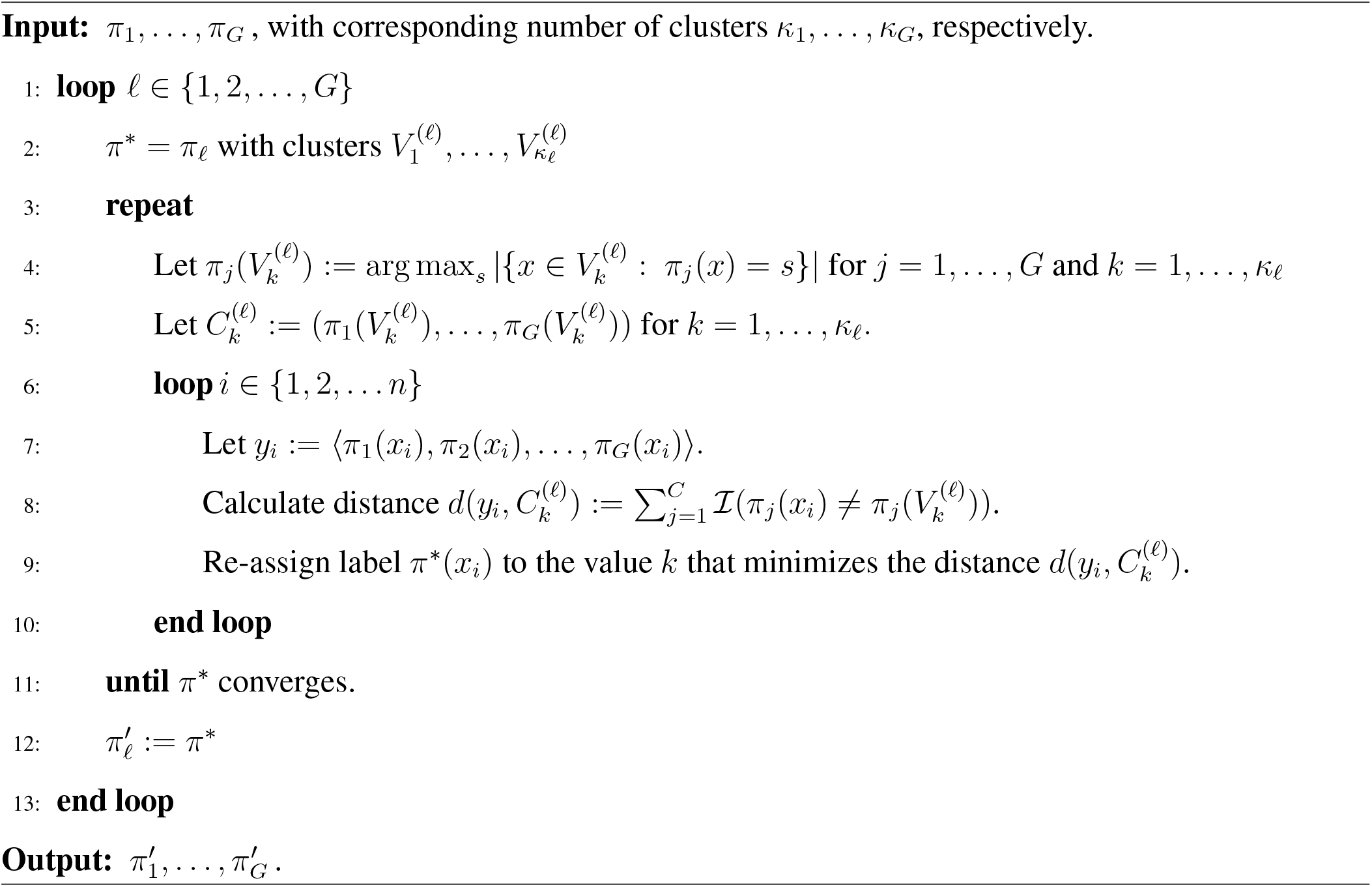

This equivalence relation defines a partition of the *n* vertices into disjoint subsets, each subset consisting of equivalent vertices. We arbitrarily number these subsets *V*_1_, …, *V*_*m*_, noting that together these are all the vertices.

A low value of *τ* results in vertices classified together in the final clustering 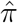 even if they are in a chain of vertices that are somewhat infrequently clustered together, while a high value of *τ* indicates that vertices are in a chain of vertices along which adjacent pairs are almost always clustered together. Therefore, in general, a high value of *τ* results in consistent inference, but may create many classes each with too few vertices.

To guarantee a minimal size for all clusters in 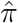 we apply a cutoff threshold *c*_*size*_ and disregard those subsets of size smaller than this threshold. The final modified iterative voting consensus clustering 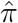 is given by {*V*_*k*_ : |*V*_*k*_| *≥ c*_*size*_}, where *c*_*size*_ is a natural number indicating the minimum cluster size. We relabel the subsets so that 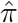 consists of 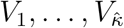 for some 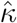. This approach does not require 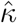 to be known a priori, nor does it require that 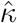 be a member of {*κ*_1_, …, *κ*_*G*_}.

Each subset *V*_*k*_ of the final clustering 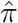 is referred to as a *connectivity-based class*. As may be observed from the clustering process, not all vertices are classified using modified IVC; vertices not paired frequently enough with other vertices are not included in the final clustering, and are eliminated from further analysis.

#### Block Probability Estimates

Finally, we estimate the block connectivity probability matrix 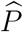 by using the clustering 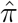 and averaging across the *G* binary matrices. Recall that 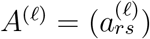corresponds to the ℓ-th binary adjacency matrix.

The *ij*-th entry of 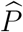 is the average proportion of connected neurons between the *i*-th and *j*-th clusters of the final clustering 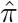, given by

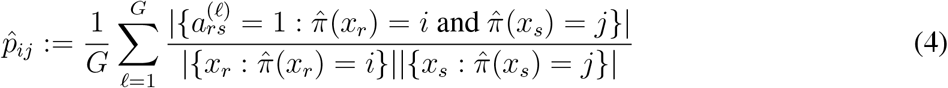

for each *r, s ∈* {1, 2, …, *n*}, and 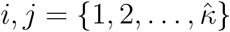. The ratio in (4) defines a value from 0 to 1.

### Statistical Analysis

#### Model Stability - Parameter Robustness

The model parameters chosen for the cluster analysis were ℙ = {*p*_conn_ = 0.15, *d* = 11, *G* = 100, *τ* = 0.95, *c*_size_ = 100}. Since the final clustering is a function of these parameters, we validate the analysis to ensure that the stochastic framework is reasonably robust with respect to the specific choice of values. Let ℙ = {*p*_conn_, *d, G, τ, c*_size_} denote the set of the five key model parameters that impact the cluster analysis, and 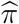 denote the final clustering (resulting from IVC) which assigns a label to every vertex. For different values of the parameters ℙ_1_, ℙ_2_, …, we obtain a corresponding clustering 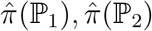 …, respectively. Unfortunately, the relation between the parameters ℙ and the clustering function 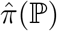 is too complex to derive an analytical expression. Also, the total number of possible combinations of valid parameter choices is extremely large, and the clustering process is computationally expensive. Therefore, it is not possible to re-perform the clustering multiple times by randomizing the choice of parameters, and use brute force to examine the clustering statistics.

Instead of randomly varying the parameters, we specifically choose only those (meaningful) parameter values which are most likely to yield a successful clustering. The set of values considered for these parameters can be found in Table 2. We compare 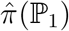 and 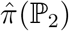 for each pair of parameters ℙ_1_ and ℙ_2_ using the *adjusted Rand index* (ARI) (Hubert & Arabie 1985). ARI is a popular similarity score for comparing two partitioning schemes for the same data points. A higher value of ARI indicates high similarity, with a (maximum) value of one indicating that they are identical and an expected value of zero for two independent clusterings.

We deem the model to be ‘stable’ if two different clusterings resulting from the same or similar parameters have an adjusted rand index of 0.90 or higher. This conservative threshold for determining a stable set of parameters ensures minimal dependence on stochastic elements of our process.

#### Comparison with other Neuronal Classification Schemes

How much of a neuron’s identity derived from network connectivity is explained by the knowledge of its neurotransmitter, or morphology (and vice versa)? To answer this question, we employ an information theoretic approach to compare and quantify the inter-dependency between different classification schemes.

Let random variable *X, Y* denote two different classifications of the same neuronal dataset, e.g., *X* represents the connectivity-based classification, and *Y* represents functional community. We then calculate the joint and conditional distributions empirically, e.g., the posterior probability *p*(*Y* = *j* | *X* = *i*) that the neuron is associated with the *j*-th functional community, given that it belongs to the *i*-th connectivity-based class. The joint and conditional distributions are in turn used to estimate the entropy *H*(*X*) (Cover 1999), which quantifies the amount of information associated with the outcome of a random variable *X*. We also estimate the mutual information *I*(*X*; *Y*), which is a measure of the mutual dependency between the two random variables. Mutual information intuitively measures how much information about *Y* is provided by knowledge of *X* alone, and vice-versa. If the two variables are independent then *I*(*X*; *Y*) = 0.

The proportion of information explained by *X* about *Y* can then be measured using normalized mutual information or NMI (Särndal 1974), often also referred to as the uncertainty coefficient (Press, Teukolsky, Vetterling, & Flannery 2007),

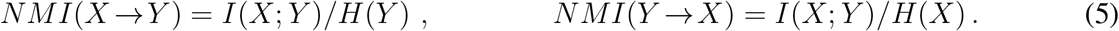

Unlike mutual information which is a symmetric measure, NMI is a directional measure while also being normalized in the [0, 1] range.

### Random Walk Model - Absorption and Driftiness

We perform a series of random walks on the derived circuit to (i) identify key nodes in the circuit, and (ii) identify important communication pathways (the most important edges in the network that connect these key nodes). The advantage of using random walks is that it provides a *dynamic* measure that takes into account redundancies in the paths, unlike neighborhood-based measures (e.g., degree centrality) or path length-based measures (e.g., betweenness centrality) which focus only on the intrinsic structure of the network. Specifically, a random walk describes a signal originating at a single node and propagating through the network via its structural connections. We refer to the origin node as the source, and the final destination node as the target. The random walk model we use to analyze connectivity patterns on our directed and weighted graph is an extension of the *absorption* and *driftiness* measures introduced in L. d. F. Costa, Batista, and Ascoli (2011).

Consider a SBM graph with *κ* number of blocks 𝒱 = {*V*_1_, *V*_2_, …, *V*_*κ*_}, and parameterized by the block connectivity probability matrix *P* = (*p*_*ij*_) *∈* [0, 1]^*κ×κ*^. The total number of vertices in the *i*-th block is denoted by |*V*_*i*_|, also referred to as the size of the block. The probability that a vertex in the *i*-th block forms an edge with a vertex in the *j*-th block, is given by an independent Bernoulli distributed with probability *p*_*ij*_. The total number of edges between any two blocks *i* and *j* is then a binomially distributed random variable, with expected value *p*_*ij*_|*V*_*i*_||*V*_*j*_|.

In our circuit representation of the SBM graph, each class corresponds to a block, and the directed, weighted edges of the circuit correspond to the block connection probabilities *P* (*V*_*i*_, *V*_*j*_) = *p*_*ij*_. A random walk on the circuit describes a sequence of *n* classes that form a path 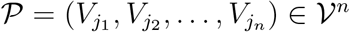. For ease of notation we re-index by setting 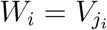 for *i* = 1, 2, …, *n*. We consider only simple paths, i.e., the random walk visits each class only once, and ignore self-loops.

Let 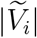 denote the scaled block sizes,

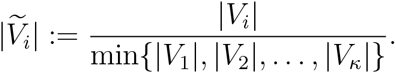

The cost of the step from class *W*_*i*_ to *W*_*i*+1_ on the path is then defined as

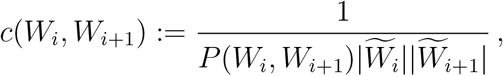

such that, the larger the expected number of edges between two blocks in the SBM graph, the lower is the cost required to traverse that particular path on the circuit.

The total path length of a random walk is obtained by summing up the cost of all steps (Supporting Information Fig. S1),

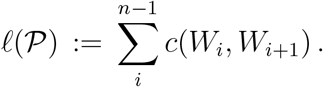

Absorption is defined as the average path length between a source class *V*_*s*_ and target class *V*_*t*_,

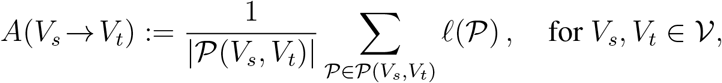

where 𝒫(*V*_*s*_, *V*_*t*_) denotes the set of all paths from *V*_*s*_ to *V*_*t*_, and |𝒫(*V*_*s*_, *V*_*t*_)| is the number of paths. The order of complexity for calculating *A*(*V*_*s*_ *→ V*_*t*_) across all possible paths and all source and target classes is larger than 𝒪(*κ*!), and is therefore computationally prohibitive for large circuits. Instead, we choose a large number *m* of random paths, which by the law of large numbers is sufficient to guarantee convergence to the true average path length. The resulting computational complexity of calculating *A*(*V*_*s*_ *→ V*_*t*_) across all source and target classes is 𝒪(*mκ*^2^).

In addition to pairwise absorption values, we also calculate the average absorption value of a random walk originating at source class *V*_*s*_ and reaching any of the other *κ −* 1 classes, and originating at any class and arriving at target class *V*_*t*_. We refer to these as average out-absorption and average in-absorption, respectively,

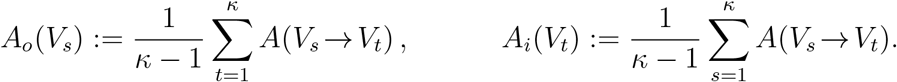

Absorption can be used to characterize important classes on the circuit. In-absorption can be used to infer the accessibility of a class. A class with low in-absorption indicates that it can be easily reached from other classes to which it connected. More accessible classes are more likely to be recruited in a wider variety of neural interactions on the circuit (L. d. F. Costa et al. 2011). On the other hand, out-absorption can be used to infer the potential that a signal originates at a class. A high out-absorption indicates that the class is located relatively upstream in the chain of signal propagation.

Driftiness is defined as the ratio between absorption and the shortest path length between the source and target node,

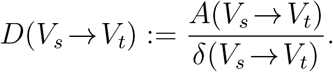

Correspondingly, the average out-driftiness and in-driftiness for each node are respectively given by

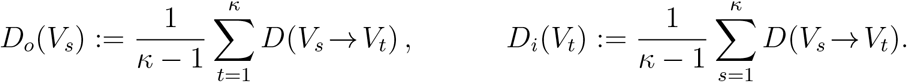

Driftiness can be used to detect important communication paths on the circuit. High driftiness indicates that the alternate paths are (on average) longer than the shortest available path, suggesting the presence of a critical individual link in the chain of signal propagation. Low driftiness indicates that the alternate paths are likely equivalent (of similar length to the shortest path), suggesting redundancy in signal propagation with multiple viable routing options.

### Periods of Critical Growth

The circuit for each day of development is obtained by only using neurons that were born on that day or before. For example, the embryo (day 0) circuit consist only of embryo neurons, whose total is denoted by *n*_0_. The day 1 circuit consists of a total of *n*_1_ number of neurons whose birthtime was either embryo or day 1, i.e., *n*_1_ = *n*_0_ + day 1 neurons. The circuit size therefore grows cumulatively, reaching on day 9 the original full adult circuit.

The circuit for the *i*-th day is a SBM parameterized by {*P*^(*i*)^, *ρ*^(*i*)^, *n*_*i*_}, where *P* ^(*i*)^ is the *κ × κ* block probability matrix, and *ρ*^(*i*)^ is the block membership probability vector 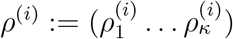, respectively.

The cluster sizes on the *i*-th day are then given by the product 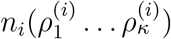 The size of the *k*-th cluster on the *i*-th day is therefore 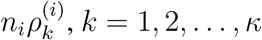 The expected reference cluster sizes are chosen 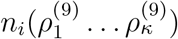. These are the cluster sizes that we would expect on *i*-th day if the final circuit was simply scaled down to *n*_*i*_. This is therefore equivalent to comparing a circuit parameterized by {*P*^(*i*)^, *ρ*^(*i*)^, *n*_*i*_} against a reference circuit parameterized by {*P* ^(*i*)^, *ρ*^(9)^, *n*_*i*_}.

Cluster growth on the *i*-th day is defined as a percentage difference between actual daily cluster growth relative to uniform growth

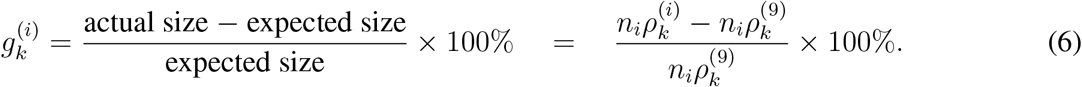

The top 25th percentile of the days which showed the greatest rate of growth as compared to the day before 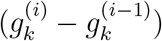, are determined to be the “critical growth period”.

### Spatial Distribution

We co-registered all 19,902 neurons to the FlyCircuit standard brain template (FCWB) (M. Costa, Manton, Ostrovsky, Prohaska, & Jefferis 2016) using the the the `natverse` R package (Bates et al. 2020). These resources are highly optimized specifically for the accurate registration of neurons from the FlyCircuit database onto 75 neuropil regions following standardized nomenclature (Ito et al. 2014), while also enabling powerful 3D visualization.

The NeuroMorpho.Org database (Ascoli, Donohue, & Halavi 2007) labels the FlyCircuit database neurons as interneurons if they only innervate adjacent neuropils, or as principal cells otherwise (Nanda et al. 2015). For each neuron, regardless of type, we derived the spatial distribution across the 75 neuropil regions by calculating the number of tracing points (re-sampled in half-micron step-sizes) within each neuropil region with the `prune_in_volume()` function of the `natverse` R package. The spatial distribution (counts and proportions of tracing points per neuropil) is provided in Supporting Information File F1 (Tab:spatial dist). Next, these 75-dimensional vectors were scaled, centered to have zero-mean and unit-variance, and orthogonalized by performing principal component analysis (PCA) (Jolliffe 2002). The orthogonalized data was then used to cluster neurons into 54 groups using the GMM-based EM algorithm (`mclust`). The resulting spatial classification (assignment of a neuron into one of the 54 groups based on the clustering) is provided in Supporting Information File F1 (Tab:all_classifications).

### Morphological Classification

We additionally performed a morphology-based classification (Bijari, Valera, López-Schier, & Ascoli 2021) by combining a selection of 15 morphological features with a quantification of branch patterns using 100-dimensional persistence homology vectors (Y. Li, Wang, Ascoli, Mitra, & Wang 2017), all obtained from NeuroMorpho.Org. The resulting 115-dimensional vectors were scaled, centered to have zero-mean and unit-variance, and then orthogonalized by PCA. Based on the elbow point (Akram, Wei, & Ascoli 2022), the first 12 principal components were used to cluster neurons into 54 groups using the GMM-based EM algorithm (`mclust`). The morphological classification (assignment of a neuron into one of the 54 groups based on the clustering) is provided in Supporting Information File F1 (Tab:all_classifications).

## RESULTS

### Mesoscale Circuit Blueprint of the Fly Brain

To construct the mesoscale level circuit we clustered the neuron-to-neuron brain-wide connectome obtained from the FlyCircuit v1.2 database (Shih et al. 2020), using the stochastic modeling framework described in subsection “Experimental Design”. Using model parameters ℙ = {*p*_conn_ = 0.15, *d* = 11, *G* = 100, *τ* = 0.95, *c*_size_ = 100}, we identified a total of 54 connectivity-based classes. Out of the 19,902 neurons, 15,571 (78.24%) were assigned to a class, while the remaining neurons were discarded from the circuit analysis. Each connectivity-based class is represented as a node in the circuit (Fig. 1a), while the directed, weighted edges represent the connection probabilities. For any two connectivity-based classes in the circuit, each neuron within one class has the same probability of forming a synaptic connection (indicated by the incident edge) with each member of the other class of neurons.

**Figure 1:**
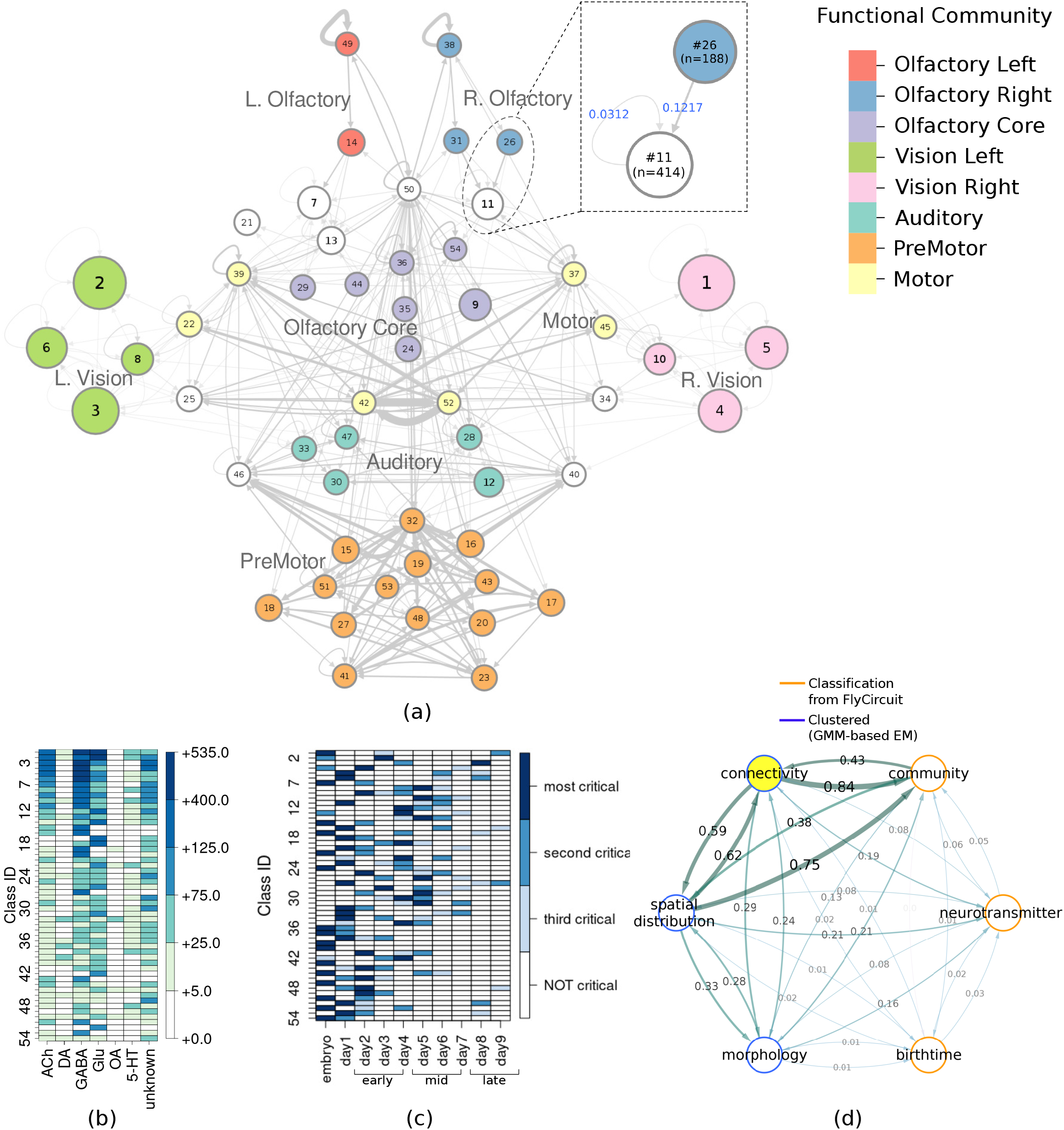
(a) Circuit diagram of 54 connectivity-based cell classes. The size of each node is proportional to the number of neurons in that cell class, while the edge thickness is proportional to the directional connection probabilities. Classes are color-coded by their dominant functional community as reported in FlyCircuit.tw (Shih et al. 2020). (b) Distribution of neurons in the 54 connectivity-based classes by neurotransmitter. ACh: acetylcholine; DA: dopamine; GABA: gamma-amino-butyric acid; Glu: glutamate; OA: octopamine; 5-HT: serotonin. (c) Mapping of connectivity-based classes by periods of critical of growth (embryo, early, mid, late), identified by the developmental birthtimes of constituent neurons. (d) Normalized mutual information among six classification schemes. A value of 1 indicates the two classifications are identical, while 0 indicates that they are independent.

The network layout views were created by using the Cytoscape software platform (Shannon et al. 2003) to organize the nodes with the prefuse force directed algorithm (Heer, Card, & Landay 2005) and grouping them by their functional community labels. The nodes were then adjusted manually based on approximate anatomical location of their constituent neuropils, while trying to maintain left-right symmetry.

By calculating the posterior conditional probabilities (as described in subsection “Comparison with other Neuronal Classification Schemes”), we identified the dominant functional community type among the neurons in a connectivity-based class, and labeled the class accordingly (Fig. 1a). Specifically, if *p*(*Y* = *j* | *X* = *i*) *>* 0.67 then the *i*-th connectivity cluster was assigned a “label” corresponding to the *j*-th functional community. Clusters with no dominant community meeting this 67% threshold criterion were not assigned any community label.

We also created similar mappings for the neurotransmitter distribution (Fig. 1b) and developmental birthtimes (Fig. 1c). These mappings enable us to characterize and interpret the circuit in relation to traditional neuronal biomarkers. For example, connectivity-based class 1 was identified as a part of right vision, contains no octopaminergic cells, and shows critical growth in late development. A detailed list of all descriptor labels assigned to each connectivity-based class, as well as the calculated prior and posterior probabilities with respect to traditional biomarkers is provided in Supporting Information File F1 (Tab:labels, Tab:prob dist). The biomarker distribution among discarded neurons broadly reflected that of the retained neurons, indicating that overall no prominent information or circuit features were missed by omitting these neurons from the analysis. For example, the prior probabilities between the discarded and the retained neurons in each Gal4 driver line had a mean absolute difference (Yitzhaki 2003) of only 0.084. Additionally, we also compiled the evolution of the circuit over the span of 10 days into a movie clip identifying critical growth periods (Supporting Information Movies M1, M2).

To ascertain how much knowledge of the traditional biomarkers is explained by neuronal connectivity and vice versa, we quantified the inter-dependency between each pair of classification schemes using normalized mutual information. The results (Fig. 1d) reveal that connectivity-based classification, neurotransmitter, and birth time are practically independent of each other (*≈* 0 *−* 0.20 NMI). We also observed very low dependency between connectivity-based classification and intrinsic morphology (*<* 0.30 NMI) indicating that connectivity-based classes may be morphologically diverse while neurons in different connectivity-based classes may share the same morphology.

While connectivity-based classification largely explains (0.84 NMI) functional communities, the reverse is not true (0.43 NMI). This is expected, as functional communities (Shih et al. 2020) were inferred by maximizing modularity (Newman 2006) to partition the network into assortative structures, characterized by denser intra-community connections and sparser inter-community connections. In contrast, SBMs not only detect assortative structures, but also disassortative and core-periphery structures (Dao, Bothorel, & Lenca 2020; Faskowitz et al. 2018; Priebe et al. 2019), providing a deeper insight into connectomic interactions (Betzel et al. 2018).

As expected, connectivity-based classification reflects certain aspects of the spatial distribution (*≈* 0.60 NMI). However, the classification obtained using only the spatial embedding of the neurons is substantially different from our connectivity-based classification, which also groups together spatially nonadjacent neurons that innervate distinct neuropils (Fig. 2a). Specifically, the SBM framework employed here groups neurons together based *exclusively* on similar patterns of stochastic connectivity, regardless of spatial location. For example, class 2 (Fig. 2b) consists predominantly (*>* 75%) of interneurons which are anatomically confined to the left lobula plate and left medulla neuropils. On the other extreme, class 50 (Fig. 2c) comprises almost entirely (*>* 99%) principal cells that are located spatially apart, and innervate a total of 21 different neuropil regions. For both these classes, the constituent neurons stably clustered together even when choosing different model parameter values (subsection “Varying the Parameters”) - including the seemingly stray neurons in class 2 which happen to have a connectivity profile similar to other fellow neurons in the class. Altogether, these results indicate that the connectivity-based classification captures latent network characteristics that are otherwise not revealed by known neuronal biomarkers.

**Figure 2:**
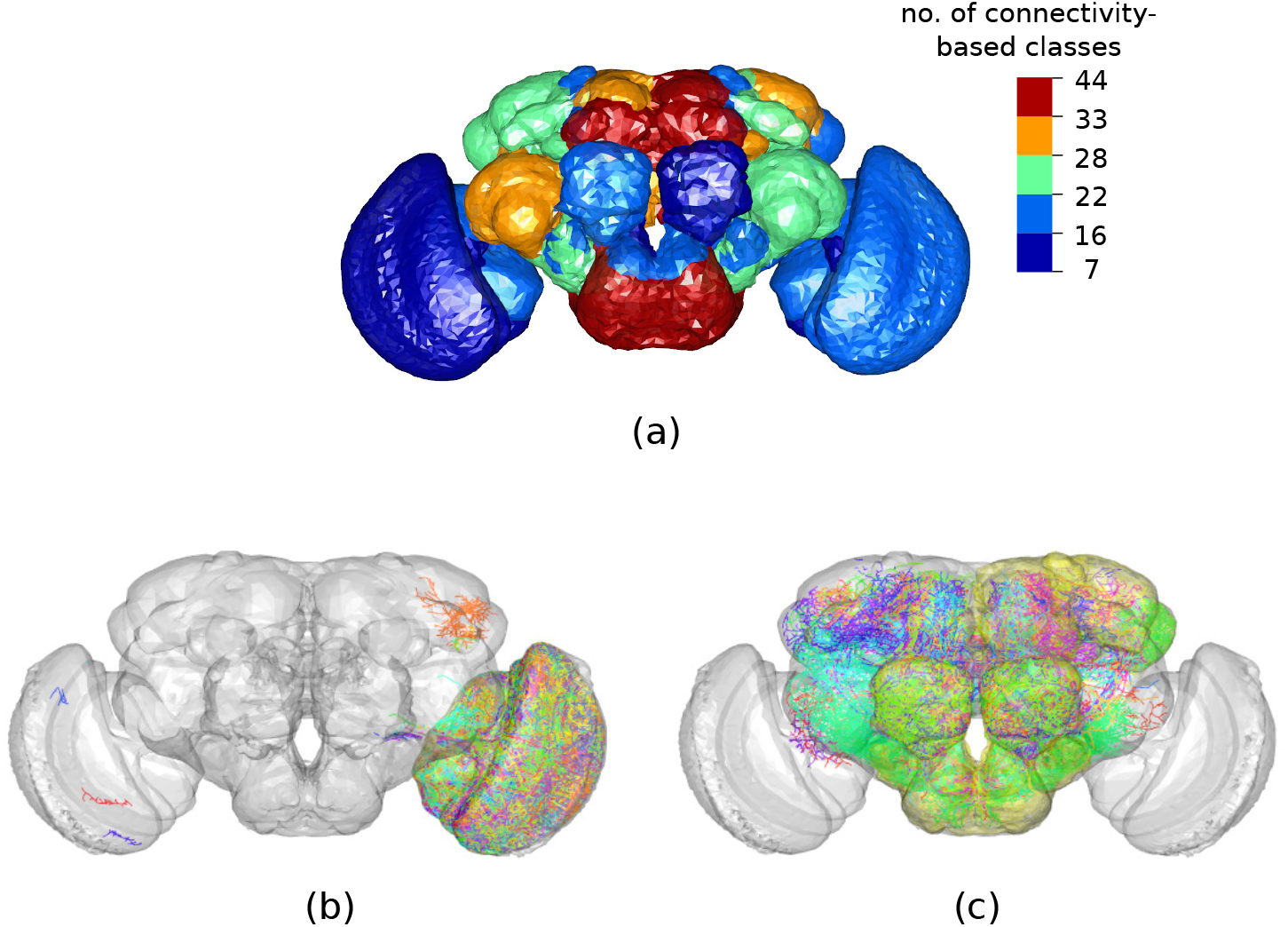
Connectivity-based classes and spatial distributions. (a) Total number of connectivity-based classes innervating each of the 75 neuropil regions, mapped using the `natverse` 3D template (Bates et al. 2020) of the *Drosophila* brain. Supporting Information Fig. S2 illustrates in detail the 3D embedding of each connectivity-based class, along with its constituent neurons and innervating neuropils. (b) Class 2, an example of spatially compact class. (c) Class 50, an example of spatially extensive class. Each neuron is assigned a random color for easier distinguishability.

### Model Validation and Stability

#### Deviation of observed data from an ideal SBM

The block probability matrix 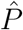 characterizing the connectivity between our identified neuronal classes was estimated using equation (4), resulting in an initial total of 1955 non-zero 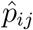’s. The median value of these 1955 non-zero entries was 0.0006. In order to better interpret the circuit, we floored all entries in 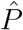 that were below a certain threshold to zero. We identified this threshold (= 0.00247) based on the largest 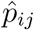 that could be set to zero while still ensuring that each class on our circuit had at least one (incoming or outgoing) edge. The estimated block connectivity probability matrix 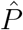 thus has a final count of 319 non-zero 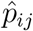’s, each representing a weighted, directed edge on our inferred mesoscale circuit (Fig. 1a). The final block connectivity probability matrix 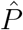 that is used to perform all subsequent analysis is provided in Supporting Information File F1 (Tab:block prob), and illustrated as a heatmap in Fig. S4a.

Recall that for a SBM, the probability that a neuron in class *V*_*i*_ forms a synaptic connection with a neuron in class *V*_*j*_ is given by an independent Bernoulli trial with probability *p*_*ij*_. If the the binary adjacency matrix *A*^(*ℓ*)^ was generated from a SBM parameterized with 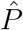, the expected number of neurons in *V*_*j*_ that any given neuron in *V*_*i*_ makes connections with would equal 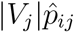 (i.e., the expected value of a binomial distribution with success probability 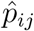, and |*V*_*j*_| trials). To assess the fit of the SBM to the observed data, we compare the observed number of connections in a block-pair (*i, j*) of *A*^(*ℓ*)^ (generated using the connectomic matrix *A*_*p*_) with the theoretical binomial distribution expected from the SBM.

Specifically, for each neuron in class *V*_*i*_ of *A*^(*ℓ*)^, we calculated the number of neurons in class *V*_*j*_ to which it is connected. We refer to this as the number of observed connections for the block-pair (*i, j*). We repeated this calculation for each of the neurons in *V*_*i*_ across all *G* = 100 matrices, giving us a total of |*V*_*i*_|*G* observation values for the block-pair (*i, j*). We then compared the spread of these observed values to the binomial distribution with parameters 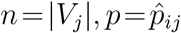. The analysis reveals that for any given block-pair the observed values are narrowly grouped around the expected value of the standard binomial, and at least 89% of the observed values lie within two standard deviations 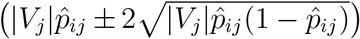.

For all 319 block-pairs that form an edge on the circuit, the percentage of observed values that lie within two standard deviations of the expected binomial mean is presented in Supporting Information Figs. S4b and S4c. Overall, these results indicate that the distribution of data in the connectomic matrices does not deviate significantly from the theoretical model predicted by the SBM.

#### Binomial Probability of Synaptic Connection

The binomial connection probability parameter *p*_conn_ was chosen to obtain a mean connection probability among potentially connected vertices of 0.5. Recall that *p*_conn_ determines the connection probability matrix *A*_*p*_, whose entries are 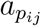 are derived from the post-synaptic connection strengths 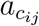 using a binomial distribution (1).

The mean connection probability *ā*_*p*_ is the mean value of non-zero entries in *A*_*p*_. We pick *p*_conn_ to be the value that minimizes |*ā*_*p*_ *−* 0.5|. Empirically, we found *p*_conn_ = 0.15.

To validate our choice of *p*_conn_, we compare our stochastically generated connectomic matrix obtained using FlyCircuit data versus the recently released Janelia hemibrain connectome (Scheffer & Meinertzhagen 2021). The edge strength of a stochastically generated FlyCircuit connectomic matrix is the number of overlapping axonal-dendritic segments that successfully form a synapse if modeled using a binomial distribution with success probability 0.15. The edge strength in the Janelia connectomic matrix is the synapse count between two neurons, experimentally determined using electron microscopy (EM). The vertex degree is the number of incoming and outgoing synapses of a neuron in the connectome (Table 1).

**Table 1:** Comparing the EM-derived Janelia hemibrain connectome (Scheffer & Meinertzhagen 2021) against our stochastically generated connectome using *p*_conn_ = 0.15. Both connectomic datasets were appropriately resized so that they span the same neuropils in the *Drosophila* hemibrain. The mean (*±* standard deviation) values for the sparsity, degree, and strength, were calculated using a total of 100 randomly sampled connectomes.

| Connectome | # Neurons | Sparsity | Avg. Vertex Deg | Avg. Edge Str |
| --- | --- | --- | --- | --- |
| EM Experimental Data (Janelia) | 21,739 | 0.75% | 326.64 | 4.04 |
| Stochastically Generated from FlyCircuit Data | 19,902 | 0.46% ± 0.01 | 181.79 ± 0.07 | 2.32 ± 0.01 |
| Truncated Connectomes: Hemibrain Only |  |  |  |  |
| EM Experimental Data (Janelia) | 11,461 | 0.75% ± 0.01 | 172.73 ± 2.23 | 4.01 ± 0.09 |
| Stochastically Generated from FlyCircuit Data | 11,461 | 0.76% ± 0.01 | 173.36 ± 0.09 | 2.58 ± 0.02 |

Since the FlyCircuit dataset has neurons sampled from the entire brain, we appropriately truncate the stochastically generated connectome by discarding all vertices that correspond to neurons that were not embedded in the neuropil regions investigated in the Janelia hemibrain dataset. We then also truncated the Janelia hemibrain connectome to match the size of the truncated FlyCircuit connectome, by randomly (uniformly) sampling neurons from the entire Janelia dataset. The sparsity and degree statistics of the resized connectomes are almost identical (Table 1), supporting our choice for *p*_conn_ = 0.15.

We also plot and compare the normalized histograms distributions (probability density functions) of the edge strength (Fig. 3a), and vertex degree (Fig. 3b) of the two truncated hemibrain connectomes. Let *f* (*x*) and *g*(*x*) denote the probability density function (PDF) of the edge strengths (or vertex degree) for the Janelia hemibrain connectome and the stochastically generated FlyCircuit connectome, respectively. To measure the similarity between the two distributions we calculate the area over which the two PDFs intersect (Cha 2007):

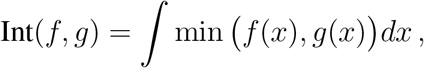

where an intersection value Int(*f, g*) = 1 indicates perfect overlap.

**Figure 3:**
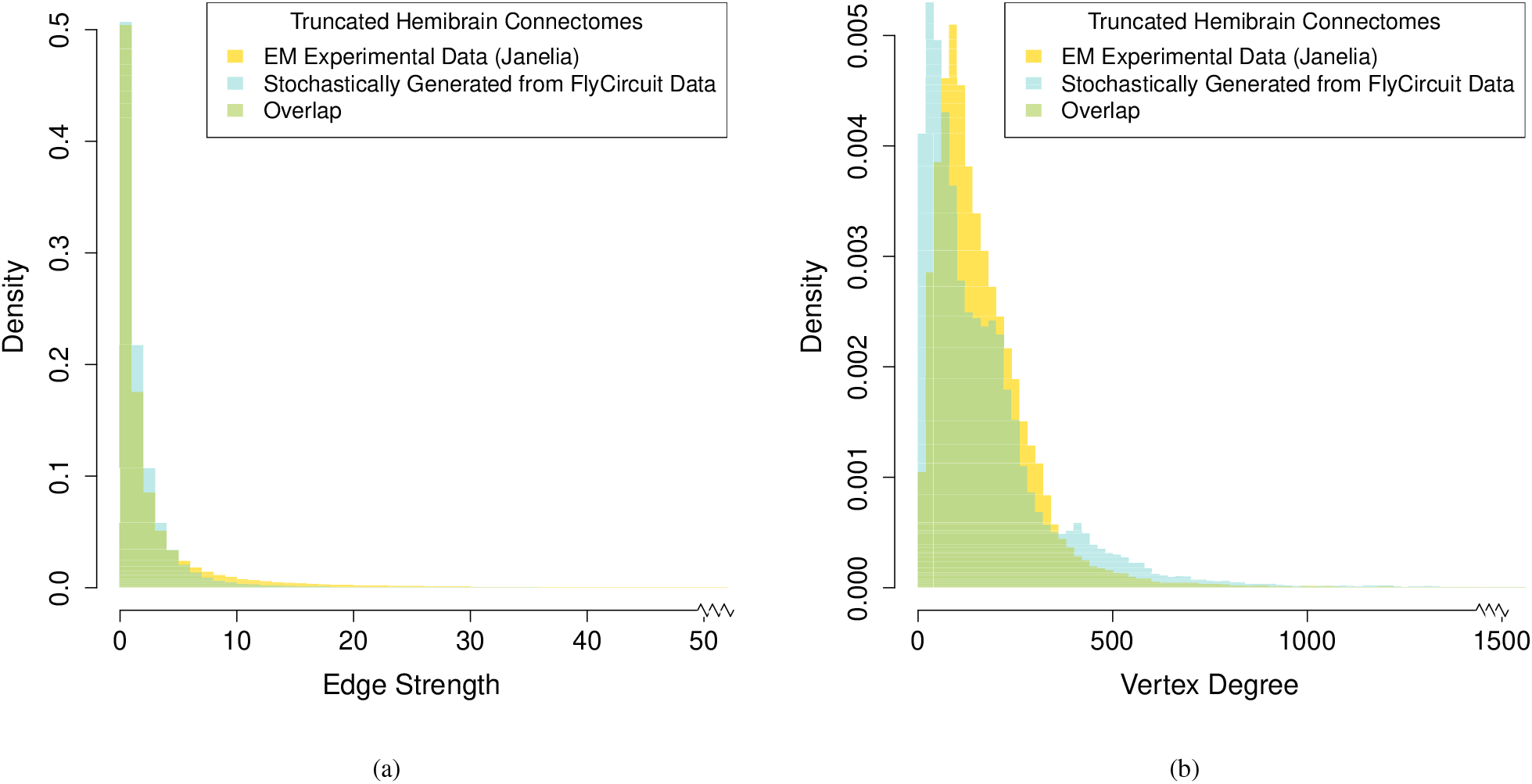
Normalized histogram distributions (probability density functions) of the (a) edge strength, and (b) vertex degree, comparing the experimentally determined Janelia hemibrain connectome (Scheffer & Meinertzhagen 2021) against our stochastically generated connectome using *p*_conn_ = 0.15. Both connectomic datasets were appropriately resized so that they span the exact same neuropils in the *Drosophila* hemibrain.

The intersection between the histogram distributions of edge strengths (Fig. 3) is 0.964, revealing that despite a difference in average edge strength due to long-tail effects, the two distributions are nearly identical. Further inspection of the Janelia experimental dataset also indicated that neurons with high- and low-edge strength in the Janelia experimental dataset appear to be evenly distributed across the neuropil regions. Moreover, the influence of very high edge strengths on our model is (exponentially) constrained as per equation (1), and for *p*_conn_ = 0.15 all edge strengths greater than 30 have *>* 0.99 probability of forming an edge in the stochastically generated binary connectome. The intersection between the histogram distributions of the vertex degree is also high with an overlap of 0.810 (Fig. 3b). While the two distributions (Fig 3b) have slightly different peaks and spread, the substantial overlap along with a nearly identical average vertex degree (Table 1) indicates that for *p*_conn_ = 0.15 the average number of edges across our multiple stochastically generated binary connectomes is comparable to the experimental dataset. The high measures of similarity between the graph-theoretical statistics of the stochastically generated connectome when compared to the EM experimental data further corroborate our choice of *p*_conn_.

#### Varying the Parameters

To validate the model stability relative to the values of all parameters in our framework (Table 2), we kept *p*_conn_ = 0.15 constant and re-performed the clustering for alternative parameter values. The possible choices for the embedding dimensionality *d ∈* {11, 15} were determined by identifying the first and second elbow-point (M. Zhu & Ghodsi 2006), respectively, on the scree plot Supporting Information (Fig. S3) of singular values. The values for the other parameters were similarly picked to yield optimal clustering (Table 2).

**Table 2:** Parameter choices for testing model stability. Values used for creating the final circuit are marked in bold.

| Parameter | Description | Value |
| --- | --- | --- |
| $p_{\text{conn}}$ | Binomial probability of synaptic connection | **0.15** |
| $d$ | Dimensionality of embedding | **11**, 15 |
| $G$ | Number of graphs | **100**, 200, 400 |
| $\tau$ | Times two neurons were placed in same cluster during IVC | 0.99, **0.95**, 0.90 |
| $c_{\text{size}}$ | Minimum cluster size | 150, 125, **100**, 75, 50 |

For example, to investigate whether increasing the number of the stochastically generated binary graphs impacted connectivity-based classification, we successively doubled only the parameter *G* while keeping all other parameters constant (bold values in Table 2). The very high pairwise adjusted rand index (ARI) values (Table 3) indicate that the resulting clusterings are all extremely similar, and that the model is stable over *G ∈* {100, 200, 400}. The percentage of neurons that are assigned a class by our framework (and not discarded by the consensus clustering and threshold parameters *τ, c*_size_) also remain similar over the range of *G*. Therefore, increasing the number of graphs beyond 100 does not substantially alter the class assignment of neurons, but only raises computational complexity. The ARI values along the diagonal entries of Table 3 were obtained by using identical parameters and number of graphs for each clustering, but repeating the stochastic generation process with different random seeds. The diagonal ARI values are therefore an indication of the dependency of the clusterings on the stochastic elements of our process, and how well we can replicate the results.

**Table 3:** Pairwise ARI values for varying the number of graphs *G ∈* {100, 200, 400} while keeping all other parameters constant {*p*_conn_ = 0.15, *d* = 11, *τ* = 0.95, *c*_size_ = 100}. The value in parenthesis indicates the percentage (out of all 19,902) neurons classified by both clusterings.

|  | 100 | 200 | 400 |
| --- | --- | --- | --- |
| 100 | 0.9514 (71.94%) | 0.9818 (72.93%) | 0.9692 (72.97%) |
| 200 | - | 0.9649 (71.92%) | 0.9737 (73.35%) |
| 400 | - | - | 0.9878 (74.89%) |

Similar ARI results for other different combinations of the parameters in Table 2 are detailed in the Supporting Information File F2. Overall, the ARI remained high (⪆ 0.90) across several different combinations, demonstrating model stability over a wide range of parameter values.

A limitation of the our inference model is the high computational cost associated with generating and then consensus clustering multiple large-scale graphs. The provided software code allows for a relatively easy parallel-processing implementation to significantly reduce the analysis time needed, but at the cost of requiring increased computational resources. The average CPU time needed for generating and clustering *G* = 100 graphs, for an embedding dimension *d* = 11 was approximately 23 hours. Doubling the number of graphs to *G* = 200 doubled the computation time, while increasing the embedding dimension to *d* = 15 only marginally increased the computation time (*≈* 24 hours). Varying *τ* and *c*_size_ had no noticeable effect on computation time. All computations were performed on a 20 core CPU cluster with Intel Haswell architecture, 64GB of RAM, and using mclust version 5.4.2.

#### Robustness to Sample Size

The connectomic dataset used in our analysis consists of 19,902 neurons, a sample corresponding to approximately 12% of the total number of neurons in a *Drosophila* brain. To investigate the robustness of our inferred circuit to the proportion of sampled neurons, we re-performed the clustering on a random subsample from the original dataset of *n* = 19,902 neurons. Each subsample consisted of a subset of *n*_*s*_ *≤ n* neurons selected using simple random sampling (without replacement). Each connectomic adjacency matrix *A*^(*ℓ*)^, for *ℓ* = 1, …, *G*, was accordingly downsized to a size *n*_*s*_*×n*_*s*_ matrix, by removing all neurons (and corresponding synaptic connections) that were not included in the subsample. The subsampled (downsized) connectomic matrices were clustered using the model parameters {*G* = 100, *p*_conn_ = 0.15, *d* = 11, *τ* = 0.95, *c*_size_ = 100*n*_*s*_*/n*}, where the threshold for minimum cluster size was adjusted to account for the reduced number of neurons in the subsample.

Table 4 shows pairwise ARI values obtained using different proportions of the original dataset, when clustering the same *G* = 100 stochastically generated connectomic matrices used to construct the circuit (Fig. 1a). The entries along the diagonal in Table 4 were obtained by comparing the clustering results for two (different) randomly selected subsamples of *n*_*s*_ neurons. The results reveal that the same analysis using 95%, or 90% of the neurons had a negligible impact on class assignment compared to using all the neurons, as the variability was comparable to repeating the stochastic generation process with different random seeds (ARI= 0.9514 for *n*_*s*_ = *n* and *G* = 100 from Table 3). Further, we observe that the number of neurons that were assigned a class by our framework (and not discarded by the consensus clustering and threshold parameters *τ, c*_size_) did not decrease even as *n*_*s*_ decreased. The classification remained fairly consistent even when using only 85% (ARI *>* 0.90) and 80% (ARI *>* 0.85) of the neurons. The fact that the class assignment of the neurons did not vary significantly for smaller proportions of the connectomic dataset makes it unlikely that our inferred brain-wide neural circuit is constricted by the sample size of the considered dataset.

**Table 4:** Pairwise ARI values comparing robustness of the clustering framework to the proportion of sampled neurons, (100*n*_*s*_*/n*)%. The value in parenthesis indicates the percentage of neurons classified by both clusterings. Note: All percentage values are expressed in relation to all 19,902 neurons.

|  | 100% | 95% | 90% | 85% | 80% |
| --- | --- | --- | --- | --- | --- |
| 100% | 1.00 (78.24%) | 0.9657 (72.18%) | 0.9424 (71.74%) | 0.9105 (72.50%) | 0.8596 (70.21%) |
| 95% | - | 0.9624 (73.67%) | 0.9516 (75.27%) | 0.9226 (74.59%) | 0.8668 (72.82%) |
| 90% | - | - | 0.9403 (75.38%) | 0.9239 (75.96%) | 0.873 (73.75%) |
| 85% | - | - | - | 0.9084 (76.00%) | 0.8779 (75.11%) |
| 80% | - | - | - | - | 0.8519 (73.08%) |

### Dopaminergic Hubs Form a Backbone Communication Pathway for Multisensory Integration

To investigate the dopaminergic pathway (Fig. 4a), we specifically consider only those neurons in each connectivity-based class whose neurotransmitter is dopamine. Recall, as per eq. (4) the connection probability 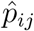 is the average proportion of connected neurons between classes *i* and *j* of the final clustering across the matrices *A*^(*ℓ*)^ for *ℓ* = 1, 2, …, 100. To identify the pathway, we used the same final clustering, but recalculated the connection probability using only the proportion of connected dopaminergic neurons in the two classes.

**Figure 4:**
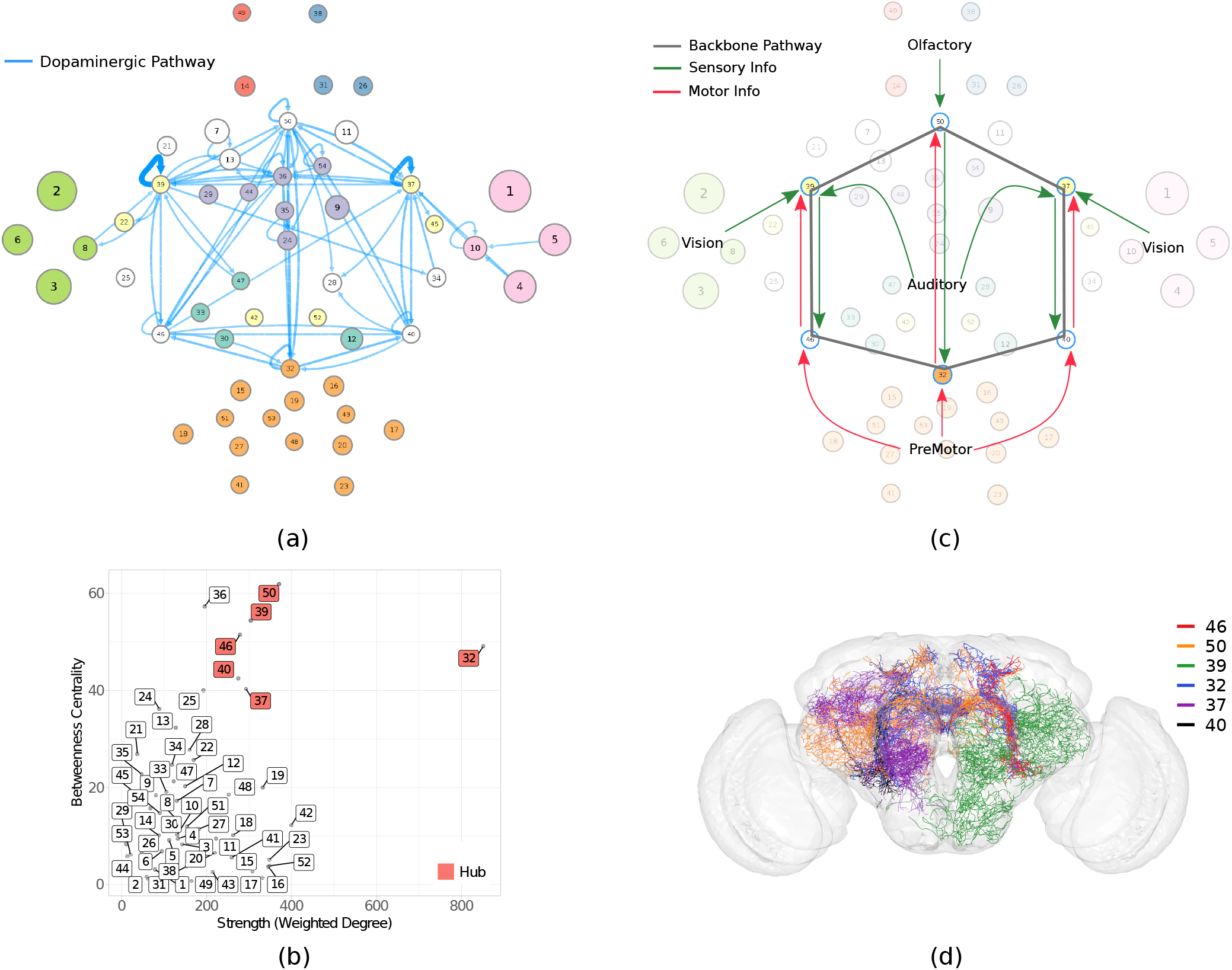
(a) Dopaminergic pathway. (b) Six classes (32, 50, 37, 39, 40, 46) have both high weighted degree and betweenness centrality, characterizing them as graph theoretical hubs in the circuit. (c) The inferred directional connectivity patterns of the identified hub classes reveal a backbone pathway for multisensory integration. The arrows represent the interpreted connectivity of the hub classes, based on important incoming and outgoing communication paths detected by analyzing the absorption and driftiness measurements for these classes (as detailed in Supporting Information Table T1). (d) Twenty-five representative dopaminergic neurons from the six hub classes that form a single path, looping over all 30 edges of the backbone communication pathway.

Six out of 54 classes (32,50,37,39,40,46) alone account for 57% (143/252) of dopaminergic neurons. These same classes also show the highest weighted degree and betweenness-centrality among all classes (Fig. 4b), characterizing them as graph theoretical hubs in the circuit. We subsequently refer to these six classes as *hub classes*. The weighted degree was defined as the sum of the incoming and outgoing connection probabilities of a class, weighted (multiplied) by the number of neurons in that class. The betweenness centrality of the class was defined by the fraction of shortest paths passing through it, where the shortest paths were calculated on a simplified circuit with unweighted edges (Brandes 2001).

The connectivity patterns on the circuit (Fig. 4c) were identified by performing a series of random walk on the entire circuit and calculating the pairwise and average absorption and driftiness values (Supporting Information File F1 (Tab:abs drift)). In general, low absorption values were used to investigate the accessibility of hub classes, while high driftiness values were used to detect the presence of important communication paths between pairs of classes (Supporting Information Table T1). For example, class 50 is the only one among all classes that is in the bottom twentieth percentile for *both* average in-absorption (seventh lowest) as well as average out-absorption (sixth lowest). These extremely low absorption values indicate that class 50 is easily accessible from other nodes on the circuit, and serves as an intermediary connector for signal propagation; thus reaffirming its role as a hub in the circuit. The connectivity arrows depicted for class 50 in Fig. 4c indicate high in-driftiness from the olfactory community, and high in- and out-driftiness from class 32, respectively. Specifically, only class 50 is in the top tenth percentile of pairwise out-driftiness values for both classes 49 and 38. The very high driftiness values between class 49 *→* 50 and 38 *→* 50 indicates that any signal originating in the antennal lobe of olfactory community is likely to arrive at class 50. Similarly, class 32 is also the only class in the top tenth percentile for both in- and out-pairwise driftiness values from class 50. We repeated this process for all six hub classes. A detailed list of *all* connectivity interpretations made on the circuit using absorption and driftiness measurements, along with a quantitative summary of these measurements, is provided in Supporting Information Table T1.

The graph theoretic analysis (Fig. 4a and 4b) and inferred connectivity patterns (Fig. 4c, Supporting Information Table T1) indicate that these six interconnected classes work together to form a backbone pathway that facilitates integration across the sensory and motor communities. The tightly interconnected loop between the hub classes observed at the circuit level was also identified at the level of individual dopaminergic neurons (Fig. 4d). Note that, although these six classes are enriched in dopaminergic neurons, dopamine is still not their dominant neurotransmitter; however, it is only the dopaminergic neurons that make these dense interconnections between the six classes. The non-dopaminergic neurons in these classes predominantly connect to sensory and motor regions. The identified pathways for all the different neurotransmitters are detailed in Supporting Information Fig. S5.

Interestingly, these hub classes also exhibit the most extensive neuronal morphologies among all classes, as characterized by highest total length and number of bifurcations. Developmental birthtimes reveal that the backbone communication pathway is one of the earliest formed pathways in the entire circuit with 90% of its connections (27/30 edges) established in embryo. The three sensory-innervated hub classes show critical growth during the embryo period, while the three premotor-innervated hub classes show critical growth on day 1.

Overall, the analysis suggests that the identified backbone pathway is critical for cross-modal integration of stimuli. The mechanics of how a fly combines input from multiple sensory modalities to guide behavior in its natural environment are not fully understood, e.g., when combining mechanosensory signals about wind direction along with visual cues to track an odor. We predict that selectively suppressing activity of neurons in the identified hub classes using the TH-Gal4 driver line will disrupt the fly’s ability to perform such complex integrative tasks.

### Glutamate-dominant Pathway bridges the Motor System inter-laterally and interlinks it with Vision to facilitate Rhythmic Activity

To identify the glutamate-dominant pathway, we recalculated the connection probabilities (4) by using only the proportion of connected glutamatergic neurons in between classes. Further, since glutamate is a common neurotransmitter in the *Drosophila* brain, we restricted our attention to the top 20% of the edges by weight (Fig. 5a), eliminating any edges below this threshold. The resultant graph theoretical pathway shows critical growth during early development, with classes 42 and 52 especially populated in embryo. These two classes also have extremely high weighted degree (but low betweenness centrality, Fig. 5b) and appear to act as a central bridge for coordinating motor activity between hemispheres.

**Figure 5:**
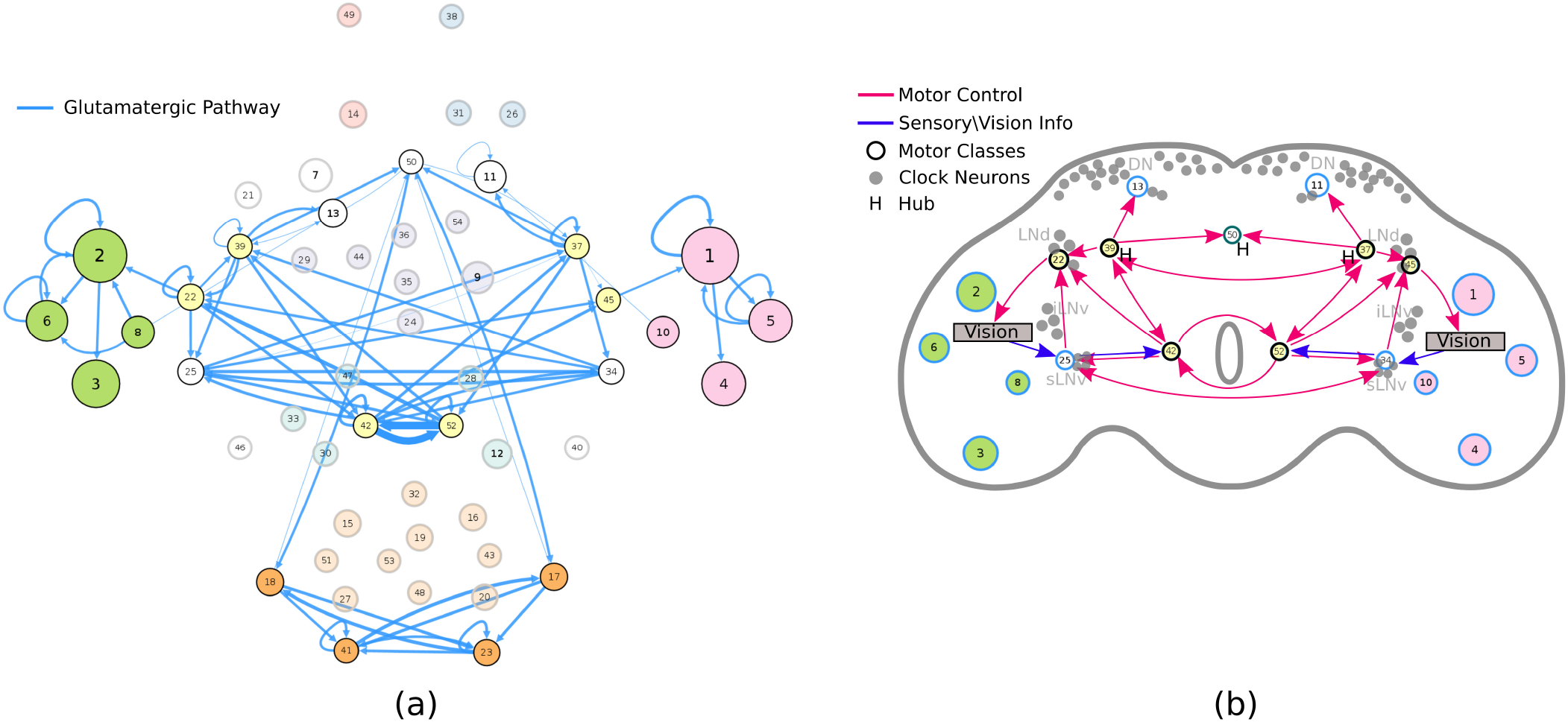
(a) Dominant glutamatergic pathway. (b) Inferred directional connectivity patterns associated with rhythmic motor activity, superimposed on a schematic of the fly brain illustrating the approximate location of the identified dominant glutamatergic classes in relation to clock neurons. The arrows represent the interpreted connectivity of the dominant glutamatergic classes, based on important incoming and outgoing communication paths detected by analyzing the absorption and driftiness measurements for these classes (as detailed in Supporting Information Table T1).

Pacemaker neurons are glutamatergic in Drosophila, implicating this neurotransmitter in behavioral rhythmicity, such as generation of locomotion and modulation of circadian photoreception (Azevedo, Hansen, Chen, Rosato, & Kyriacou 2020; Guo et al. 2016; Zimmerman et al. 2017). Although the anatomical locations of clock neurons in the Drosophila brain are known, their connectivity pathways remain largely unexplored (Dubnau 2014). Circadian properties may be an emergent function of the the underlying circuitry and not the property of clock neurons alone (Azevedo et al. 2020; Dissel et al. 2014).

Random walk analysis over the circuit (Fig. 5b) revealed that classes 22 and 25 in the left hemisphere, and 34 and 45 in the right hemisphere, serve as the primary gateways between the vision and motor systems (Supporting Information Table T1). Specifically, these classes receive signals from the visual system and pass on this information to the downstream motor regions, while also transmitting motor signals back to vision – possibly control signals to modulate input. Further, the inferred graph theoretical pathway features multiple inter-connecting loops in the motor region, resembling feedback control, with strong interaction between left and right hemispheres. We predict that suppressing the activity of glutamatergic neurons in these identified classes, but not elsewhere, by VGlut-GAL4 driver line targeting will disrupt motor patterning and circadian rhythms.

### Mechanosensory Pathways rely on Parallel Serotonergic and Octopaminergic Neurotransmission

We identified connectivity-based classes associated with sensory-motor processing through mappings to functional communities and spatial distribution. Specifically, we examined those classes whose neurons heavily innervate the central complex and are identified to be part of the premotor, auditory, or motor communities. The central complex has been linked to spatial representation, spatial orientation, visual place learning, and navigation (Turner-Evans & Jayaraman 2016). These combined pathways are also believed to be associated with other mechanosensory functions such as gravity sensation, wind sensation, and air current feedback during flight (Boekhoff-Falk & Eberl 2014).

The serotonergic pathway (Fig. 6a) was identified by restricting attention to only the classes identified above, and recalculating the connection probabilities (4) using only the proportion of connected serotonergic neurons in these classes. The serotonergic system is a critical component of place learning (Zars 2009). The octopaminergic pathway was similarly identified by considering only the octopaminergic neurons in the same classes as above (Fig. 6b). While there are only 140 octopaminergic neurons in our connectome, octopamine is critical for essentially all sensory inputs and is known to play a role in fight-or-flight response (Sotnikova & Gainetdinov 2009), stress-related enhancement of motor activity (Sujkowski, Ramesh, Brockmann, & Wessells 2017), and aggression (Hoyer et al. 2008). Compared to the premotor region which shows critical growth in the early period, the auditory classes grow predominantly during mid-development, with the octopaminergic classes (19, 20, and 27) showing continued growth even in the latest stages. Interestingly, while running in parallel throughout most of the mechanosensory system, the two graph theoretical pathways diverge along the dorsal-ventral axis (Fig. 6c), with octopaminergic neurons predominantly innervating the gnathal ganglion and ventromedial protocerebrum, and serotonergic neurons innervating the superior lateral protocerebrum and superior medial protocerebrum.

**Figure 6:**
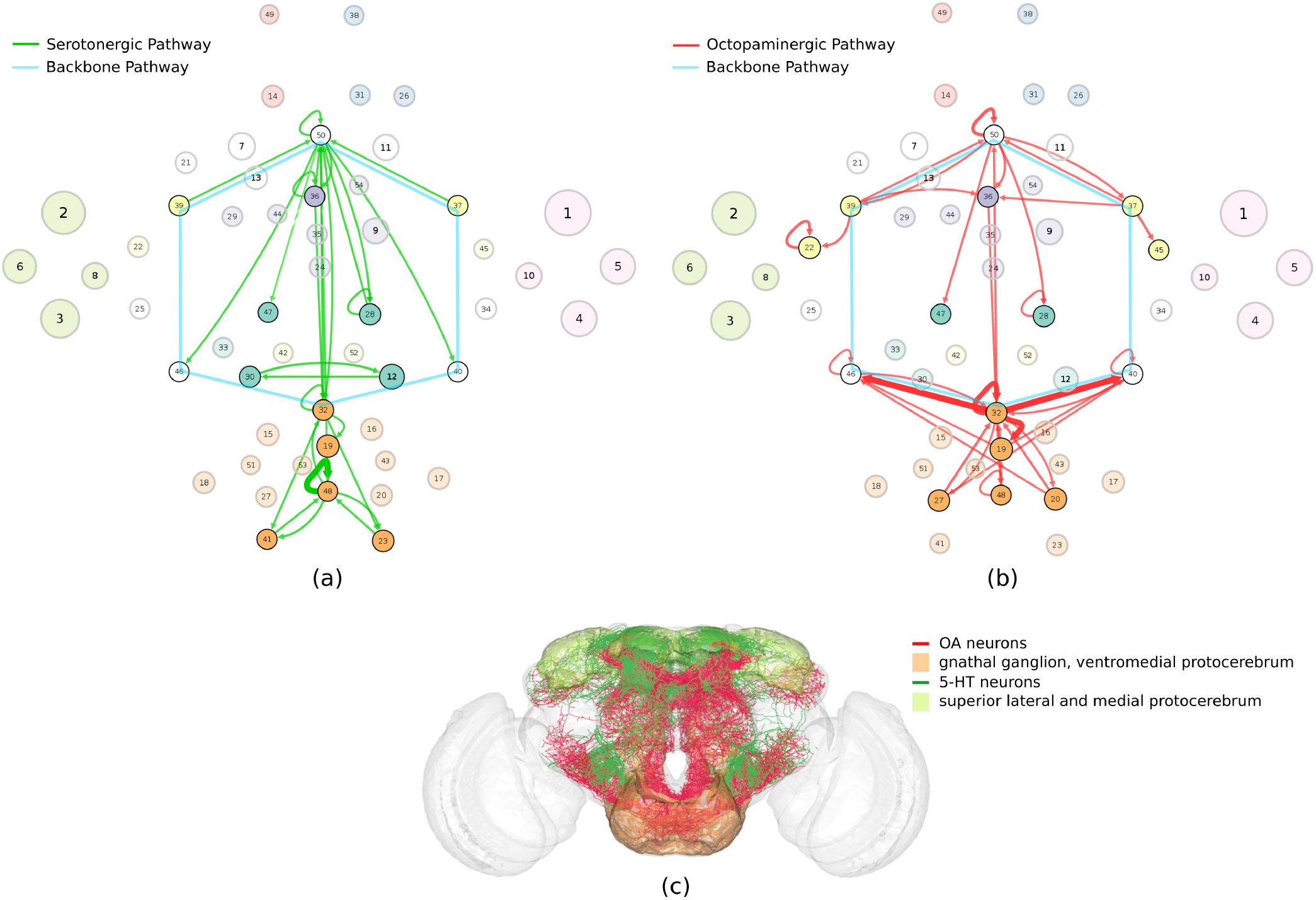
(a) Serotonergic pathway associated with spatial orientation. (b) Octopaminergic pathway associated with fight-or-flight response. (c) The mechanosensory pathways diverge along the dorsal-ventral axis with serotonergic and octopaminergic neurons innervating distinct neuropils.

The analysis suggests that these parallel pathways aid in navigation and place learning by working in conjunction with the hub classes on the backbone pathway to gather information using sensory cues during flight. We predict that inhibiting the serotonergic neurons in this pathway using the Trh-Gal4 driver would interfere with the fly’s ability to orient itself in relation to its surrounding. In addition, we hypothesize that selectively inhibiting octopaminergic neurons in this pathway using the Tdc2-Gal4 driver would prevent the fly from responding appropriately to potential threats such as moving away from a predator.

### Olfactory Learning depends on non-redundant GABAergic and Glutamatergic signaling, while Odor Tracking relies on fast Serotonergic Pathways

To identify the circuit pathways involved in olfaction, we restricted our attention to those classes that are associated with the olfactory communities, and whose neurons heavily innervate the antennal lobes, lateral horn, and mushroom body. The mushroom body plays a central role in associative learning and memory, particularly of olfactory information (Busto, Cervantes-Sandoval, & Davis 2010). Within the above identified classes, we considered the neurons associated with the three neurotransmitters involved in olfactory processing, namely GABA (Busto et al. 2010), glutamate (Liu & Wilson 2013), and serotonin (Johnson, Becnel, & Nichols 2011). The olfactory pathways (Fig. 7) were identified by recalculating the connection probabilities using only the proportion of connected GABA (or glutamatergic, or serotonergic neurons) in these specific classes. The identified graph theoretical pathways are consistent with findings that GABA and glutamate function in parallel to gate olfactory input from the antennal lobe to the mushroom body (Liu & Wilson 2013).

**Figure 7:**
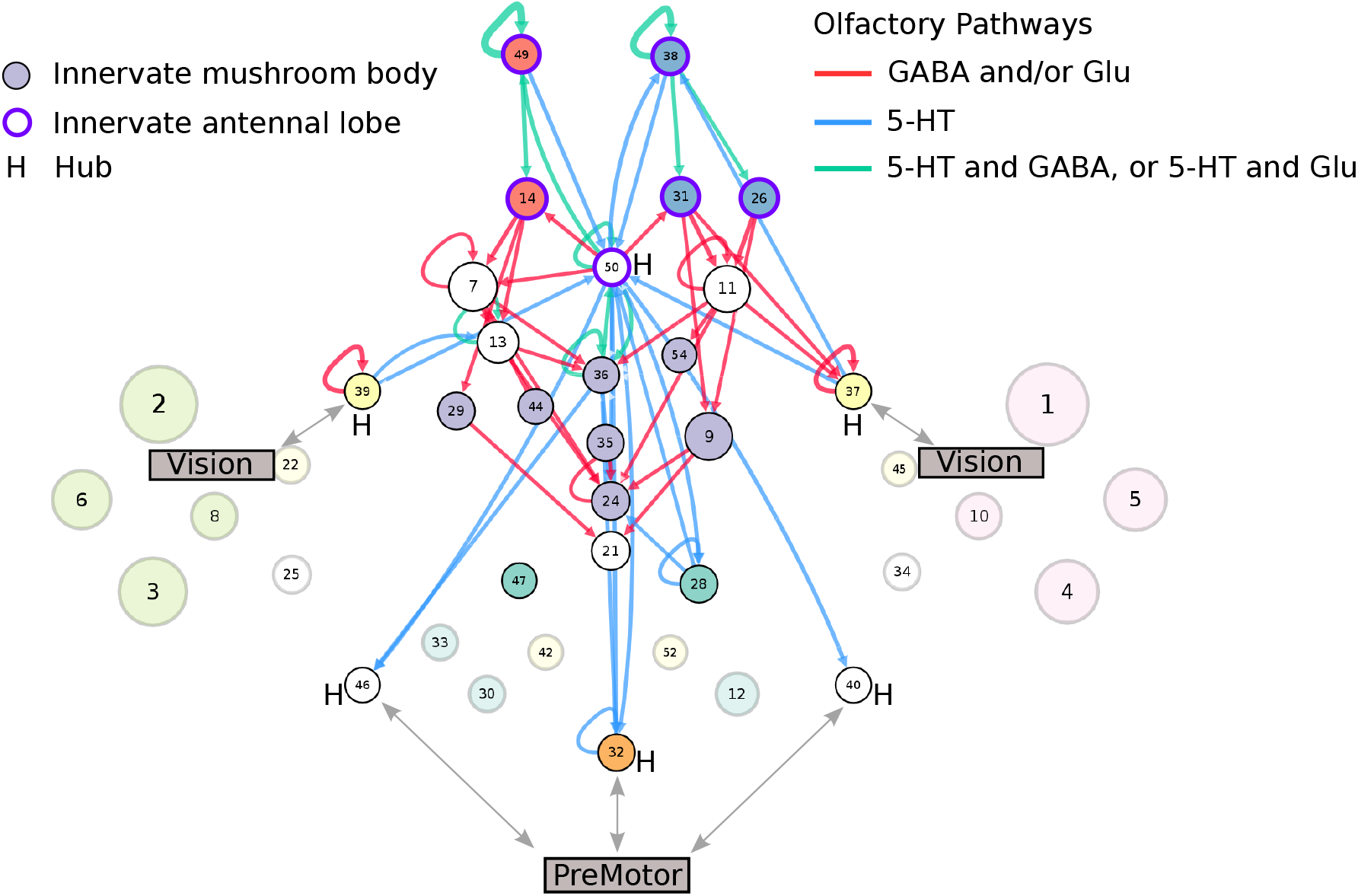
Predicted olfactory pathways. GABA and glutamate function in parallel to enable olfactory learning, while serotonergic connections enable cross-modal integration to drive odor tracking.

Random walk analysis revealed that these pathways have predominantly short local connections with important individual paths and uncharacteristically few alternative communication routes. Thus, we predict that selectively suppressing glutamatergic (via VGlut-Gal4 driver) and GABAergic (via Gad1-Gal4) transmission in this pathway would impact olfactory learning tasks such as discriminating among similar odors or varying odor concentrations. Since VGluT-Gal4 and Gad1-Gal4 cover thousands of glutamatergic and GABAergic neurons, respectively, throughout the fly brain, selectively blocking neurotransmission in this circuit could harness split Gal4 lines (Luan, Diao, Scott, & White 2020) or other approaches to spatially target neurons specifically in the olfactory system (Zanon, Zanini, & Haase 2022).

Flies are known to track odor by integrating cross-modal mechanosensory and visual input in flight (Duistermars & Frye 2010), yet little is known about these descending neural pathways. While the inferred graph theoretical pathways reveal strong olfaction-premotor and olfaction-vision link via the hub classes, we discovered that there are very few serotonergic connections to or from the mushroom body (olfactory core) classes themselves. Birthtime analysis shows that the critical growth period of the olfaction-vision connections occurs in early development, while the olfaction-premotor connections are already present in embryo. We hypothesize that this pathway enables fast connections from a stimulus (antennal lobe) to a behavioral response (premotor), and that selective blockage of those serotonergic neurons via Trh-Gal4 driver line targeting would interfere with odor localization.

## DISCUSSION

Over the last two decades, *Drosophila* has been a popular model organism for studying how structural connections in the brain give rise to functional interactions. Numerous studies have previously focused on the functional dissection of *regional* neural circuitry in *Drosophila*, including that of vision (Borst, Haag, & Mauss 2020; Y. Zhu 2013), olfaction (Busto et al. 2010), mushroom body (F. Li et al. 2020), and central complex (Turner-Evans & Jayaraman 2016), to name a few. With advancements in data acquisition and processing, recent large-scale projects (Scheffer & Meinertzhagen 2021; Shih et al. 2015) have been successful in generating detailed connectomes of the *brain-wide* neural circuitry, that go well beyond a single region. While a complete brain-wide wiring diagram is a necessary prerequisite, unraveling the underlying mechanistic pathways to interpolate function and behavior is a problem that cannot be solved by gathering more data alone (Scheffer & Meinertzhagen 2021). One of the fundamental obstacles is the absence of a quantitative specification of neuron types and how they relate to neural circuitry.

The prevalent approach to analyzing brain-wide microscopic connectomes is to group neurons into spatially compact regions, e.g., local processing units (Shih et al. 2015) or compartments (Scheffer et al. 2020), and then attempt to characterize the connectivity between these groups. While there is undoubtedly a strong spatial component to synaptic connectivity, anatomical division is by itself insufficient to deconstruct brain computation. Modeling connectomes as a network comprising solely of assortative, anatomically segregated regions interacting with each other (Shih et al. 2020) may crucially miss many functionally relevant characteristics pertaining to the complex mesoscale organization of the brain.

In contrast, our SBM framework groups neurons by their patterns of potential synaptic connectivity to capture assortative and non-assortative features in the underlying neuron-to-neuron brain-wide network. The information captured by connectivity-based classification is complementary, yet synergistic, to that provided by traditional neuronal classification - together enabling functional interpretation not previously possible. By inferring the directed connectivity patterns on the derived circuit and simultaneously characterizing the circuit with respect to multiple biomarkers, we have identified functional pathways supporting multiple sensory modalities and cross-region integration.

A key pathway identified by our circuit is the backbone communication pathway comprising of six interconnected dopaminergic-centric hubs working together to facilitate integration of the sensory and motor pathways. The neural circuitry for vision, olfaction, and hearing are well characterized, but the pathways for how these modalities combine to guide behavior (Currier & Nagel 2020), e.g., during navigation, remain elusive. Interestingly, two out of these six hubs are the only classes overall that consist of all three neuromodulators — DA, OA, and 5-HT. Further, since the hubs innervated multiple neuropils, this pathway would not have been revealed by a classification purely based on anatomy, neurotransmitter, or morphology. Dopamine is a known neuromodulator central to reinforcement signaling (Felsenberg, Barnstedt, Cognigni, Lin, & Waddell 2017; Waddell 2013), and responding to salient stimuli related to olfaction and vision (Kasture, Hummel, Sucic, & Freissmuth 2018). Also, while dopamine affects visual tracking (Riemensperger et al. 2011), its role in place learning or spatial memory remains to be elucidated (Zars 2009).

Another prominent pathway revealed in our analysis is the glutamate-dominant pathway for circadian activity. The clock neuron subsets in the *Drosophila* brain are well described including their spatial location, morphology, neurotransmitter, and role in behavioral rhythmicity (Azevedo et al. 2020; Guo et al. 2016; Zimmerman et al. 2017). Nevertheless, the inputs from other neurons onto clock neurons, and the exact neuronal pathways underlying the entrainment of the clock for synchronization with the environment are poorly known (Azevedo et al. 2020; Dubnau 2014; Hamasaka, Wegener, & Nässel 2005). The identified pathways link the motor and vision regions inter-laterally, suggesting involvement in coordination of rhythmic activity. Unfortunately, however, it is unknown which, if any, of the neurons in the 19,902-neuron FlyCircuit dataset are actually clock neurons.

Over the last several years, optogenetics (Kohsaka & Nose 2021) have been invaluable in studying how dynamical interactions on the neural circuit give rise to specific *Drosophila* behavior. These include circuits relating to rhythmic motor function (Ravbar, Zhang, & Simpson 2021), olfaction (Bellmann et al. 2010; Hige, Aso, Modi, Rubin, & Turner 2015; Owald et al. 2015), central complex function (Franconville, Beron, & Jayaraman 2018), and learning and memory (Simpson & Looger 2018). Our derived circuit is amenable to optogenetic manipulation, thus allowing for experimental verification of the predicted functional behavior of the identified pathways. Specifically, for each of the characterized circuit pathways, we identify GAL4-UAS driver lines (Brand & Perrimon 1993) that could potentially be used for selective neuronal targeting. We also predict the associated loss (or gain) of function that would accompany the selective deactivation (or activation) of the corresponding neurons in these pathways.

The connectomic dataset used in our analysis consists of 19,902 neurons reconstructed from multiple specimen using light microscopy, corresponding to a sample size of approximately 12% of the total number of neurons in a *Drosophila* brain. As with all research based on sampled data, we cannot exclude the possibility that our analysis may miss some essential features of the brain-wide circuitry, due to important neural classes being absent or not represented with enough neurons in the considered dataset. Our derived mesoscale circuit does however demonstrate robustness to the proportion of neurons sampled, suggesting that small variations in the size of the dataset are unlikely to change conclusions significantly. The constructed circuit additionally shows excellent left-right symmetry, indicating bilaterally uniform sampling of the neurons.

A practical limitation of applying the spectral graph clustering framework is that it requires the size of the dataset (number of vertices in the network) to be much larger than the number of blocks/classes *κ*. In particular, *κ << n* may not hold for connectomic datasets with a limited number of sampled neurons, or with a large number of classes in relation to total neurons. On the other hand, the SBM inference model scales extremely well as *n* increases (Mehta et al. 2021; Yang, Priebe, Park, & Marchette 2020); thus, this analysis can be progressively refined as more neurons continue to be morphologically reconstructed, registered to a common atlas, and shared with the community (Ascoli, Maraver, Nanda, Polavaram, & Armañanzas 2017);.

Our model could also be applied to dense connectomic reconstructions from electron microscopy, which is considered the gold standard in the field. However, we view the knowledge derived from sparse light microscopy collected from multiple brains valuable in its own merit, as it provides complementary information on the shared circuit elements across individuals (Ascoli 2015). In this regard, it is useful to note that our probabilistic model, currently restricted to Bernoulli adjacency matrices, can be generalized to cluster non-binary connectomes using a weighted SBM (Aicher, Jacobs, & Clauset 2015; Ng & Murphy 2021), which would allow accounting for the number of synapses between connected neuron pairs (Tecuatl, Wheeler, & Ascoli 2021) in the identification of connectivity-based classes.

## DATA AND CODE AVAILABILITY

All the data, code, and relevant information to replicate the analysis and results described in this article are made publicly available at github.com/k3t3n/FlyConn (Mehta, Goldin, & Ascoli 2022).

## SUPPORTING INFORMATION

Supporting information for this article (also available at github.com/k3t3n/FlyConn) includes the following supplementary material. Figure S1: Random walk path length. Figure S2: Scree plot. Figure S3: 3D spatial embedding of each class. Figure S4: Block connectivity probability matrix.

Figure S5: Neurotransmitter pathways. Table T1: Absorption and driftiness. File F1: Class labels and data. File F2: ARI results. Movie M1: Critical growth per day. Movie M2: Critical growth time periods.

## ACKNOWLEDGMENTS

We thank our colleagues at the Center for Neural Informatics, Structures, and Plasticity (CN3) for many insightful discussions. We are especially grateful to Dr. Sumit Nanda for feedback on the neuropil distribution of the FlyCircuit reconstructions and to Dr. Carolina Tecuatl for help with the Janelia hemibrain connectome tracing files.

