## Supplementary material for "Circuit Analysis of the Drosophila Brain using Connectivity-based Neuronal Classification Reveals Organization of Key Communication Pathways": Figure S1.pdf

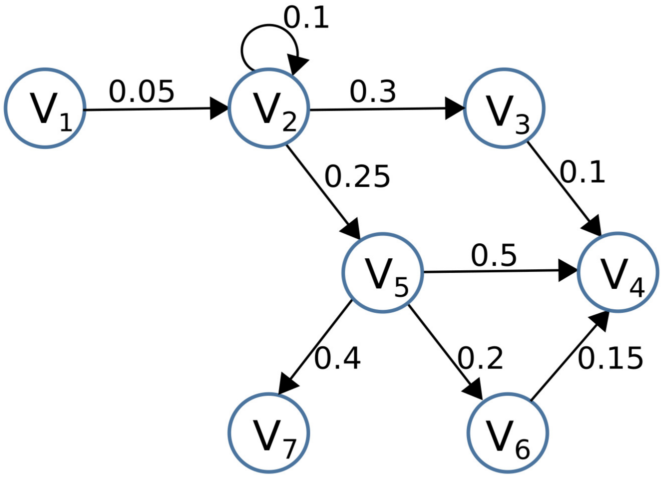


Number of neurons in *i*-th class

| $i$ | $\lvert V_i \rvert$ | $\lvert \tilde{V}_i \rvert$ |
|---|---|---|
| 1 | 1000 | 10 |
| 2 | 250 | 2.5 |
| 3 | 100 | 1 |
| 4 | 500 | 5 |
| 5 | 300 | 3 |
| 6 | 400 | 4 |
| 7 | 150 | 1.5 |

Block connection prob. $p_{ij}$

| | | | | | | |
|---|---|---|---|---|---|---|
| 0 | 0.05 | 0 | 0 | 0 | 0 | 0 |
| 0 | 0.1 | 0.3 | 0 | 0.25 | 0 | 0 |
| 0 | 0 | 0 | 0.1 | 0 | 0 | 0 |
| 0 | 0 | 0 | 0 | 0 | 0 | 0 |
| 0 | 0 | 0 | 0.5 | 0 | 0.2 | 0.4 |
| 0 | 0 | 0 | 0.15 | 0 | 0 | 0 |
| 0 | 0 | 0 | 0 | 0 | 0 | 0 |

Cost $c_{ij} := \dfrac{1}{p_{ij}|\widetilde{V}_i||\widetilde{V}_j|}$

| | | | | | | |
|---|---|---|---|---|---|---|
| 0 | 0.8 | 0 | 0 | 0 | 0 | 0 |
| 0 | 0 | 1.33 | 0 | 0.53 | 0 | 0 |
| 0 | 0 | 0 | 2 | 0 | 0 | 0 |
| 0 | 0 | 0 | 0 | 0 | 0 | 0 |
| 0 | 0 | 0 | 0.13 | 0 | 0.42 | 0.56 |
| 0 | 0 | 0 | 0.33 | 0 | 0 | 0 |
| 0 | 0 | 0 | 0 | 0 | 0 | 0 |

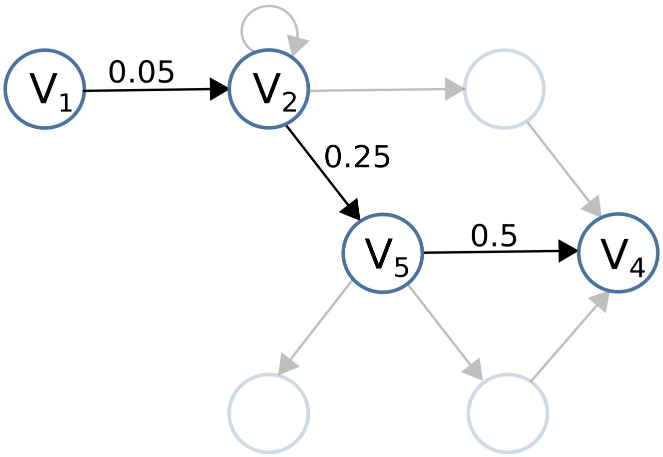


Path:

$$\mathcal{P} = \{V_1, V_2, V_5, V_4\}$$

Path length:

$$\ell(\mathcal{P}) = c_{12} + c_{25} + c_{54}$$

$$= 0.8 + 0.53 + 0.42$$

$$= 1.75$$

**Figure S1:** A simple example illustrating how the path length of a random walk is calculated, using a mock circuit with seven classes (blocks). The number of neurons in each class $|V_i|$, and directional edge weights between classes $p_{ij}$ are assumed to be known — which are used to calculate the cost $c_{ij}$ of each random step in the walk. The path length for the random walk $V_1 \to V_2 \to V_5 \to V_4$ is then obtained by summing the costs along the traversed path.
