## Supplementary material for "Circuit Analysis of the Drosophila Brain using Connectivity-based Neuronal Classification Reveals Organization of Key Communication Pathways": Figure S2.pdf

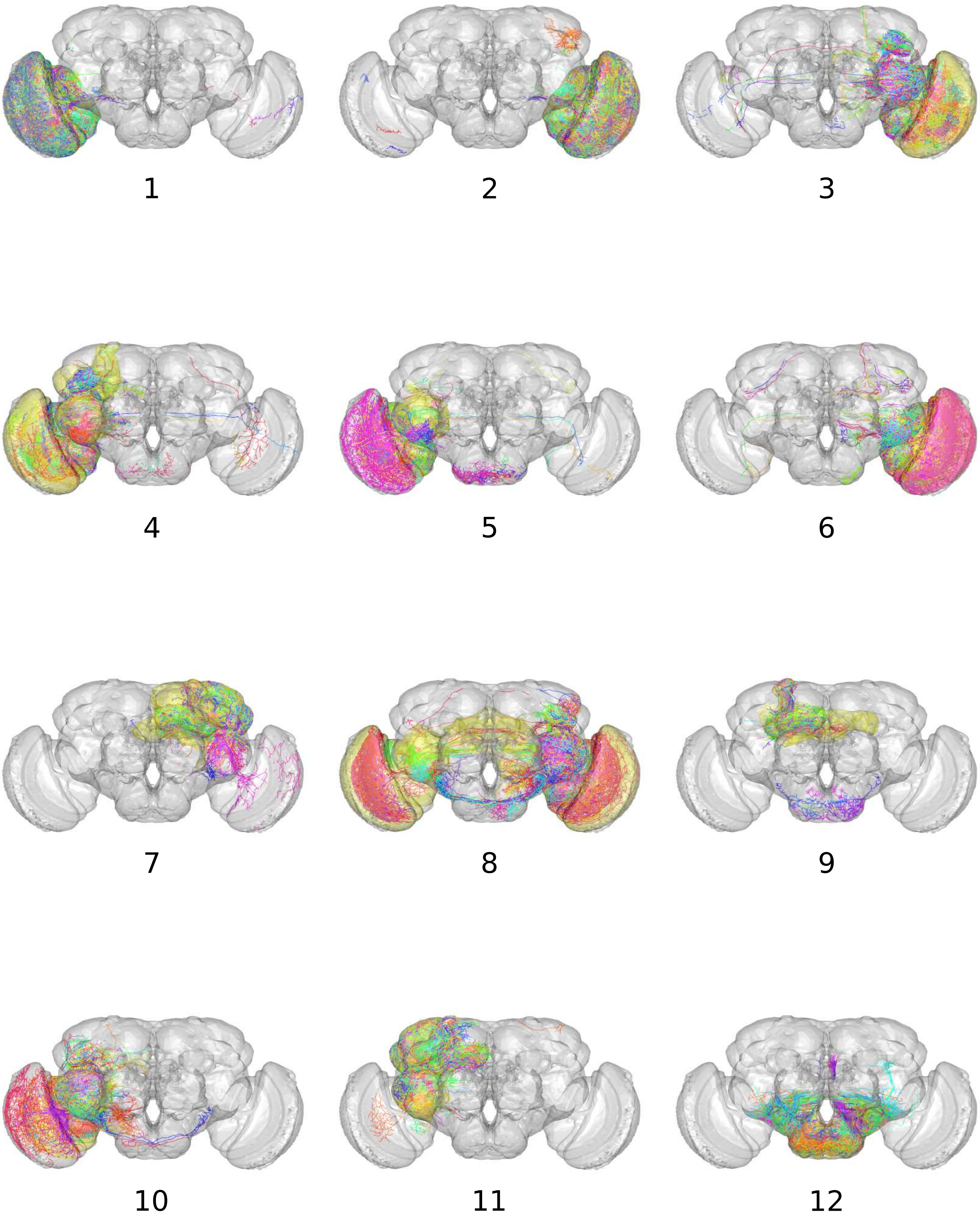

1
2
3
4
5
6
7
8
9
10
11
12

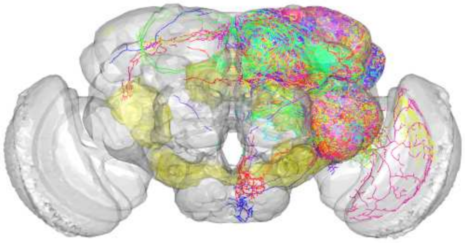

13

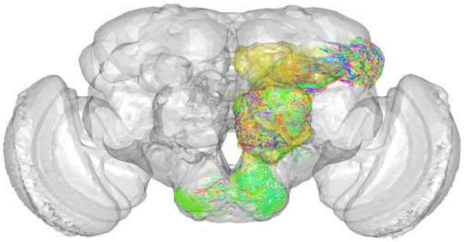

14

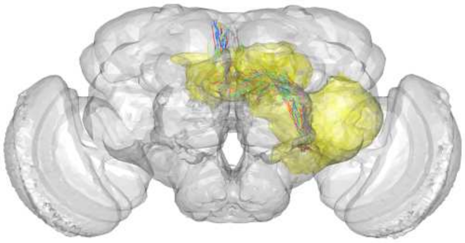

15

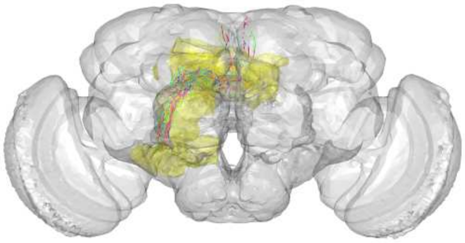

16

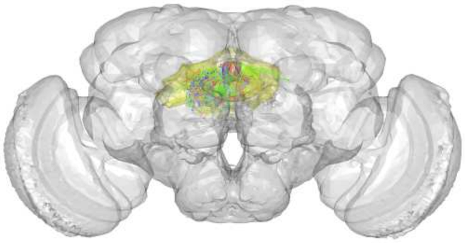

17

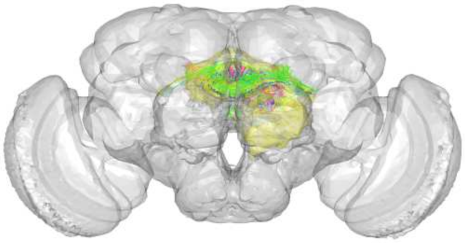

18

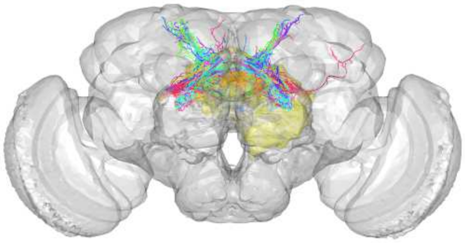

19

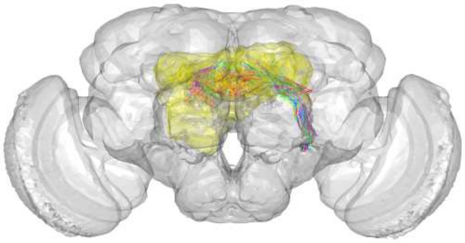

20

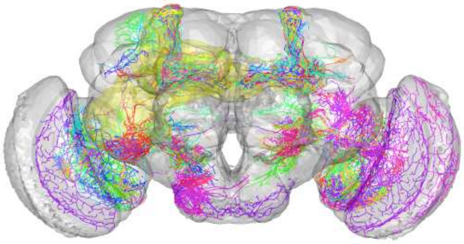

21

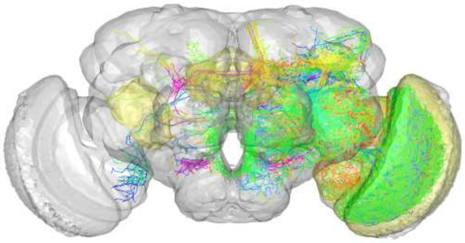

22

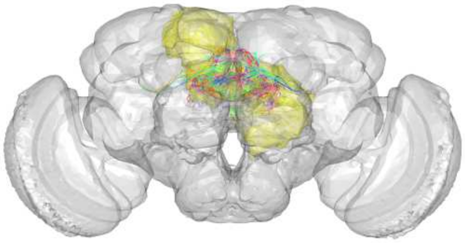

23

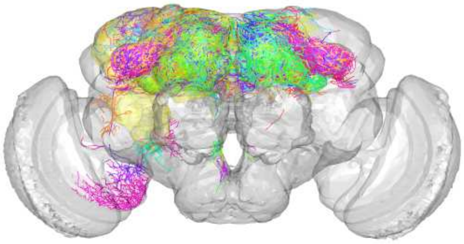

24

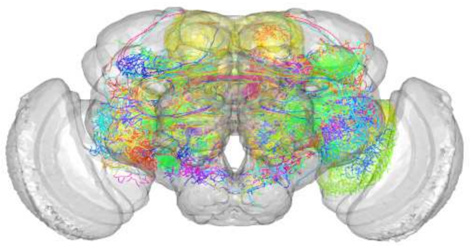

25

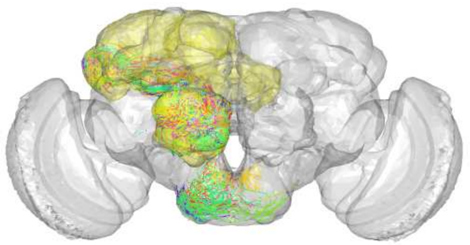

26

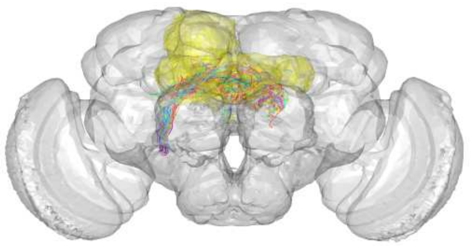

27

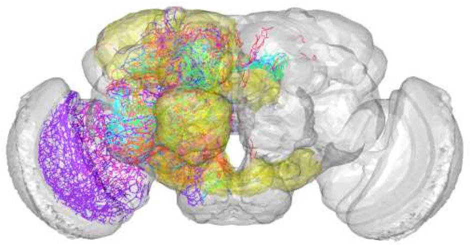

28

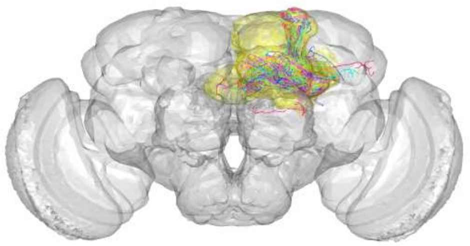

29

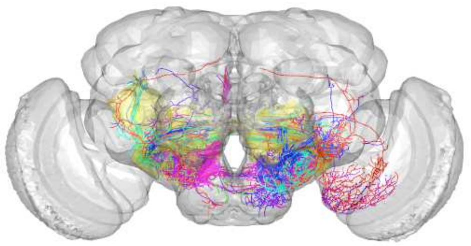

30

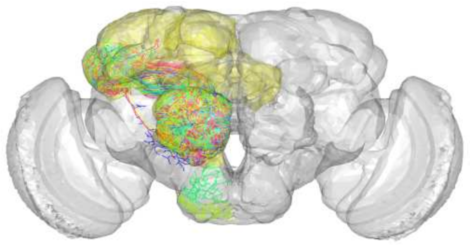

31

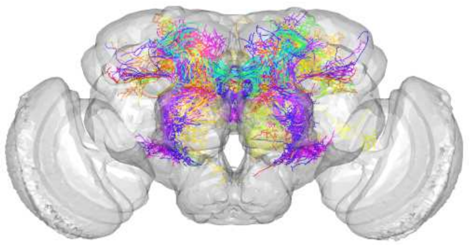

32

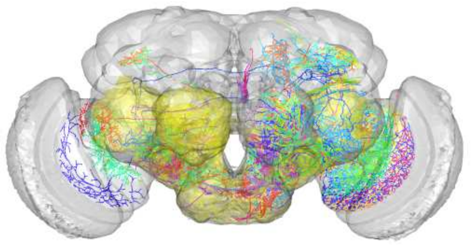

33

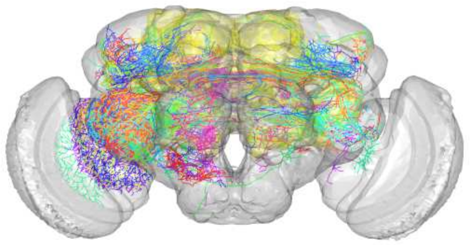

34

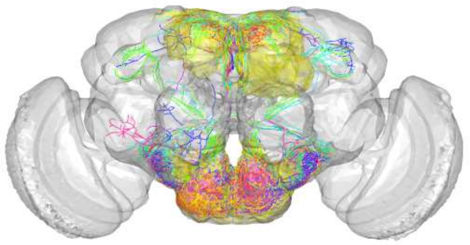

35

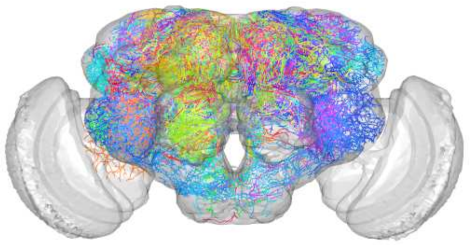

36

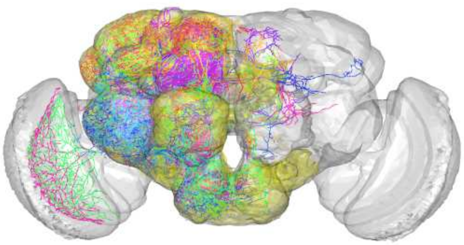

37

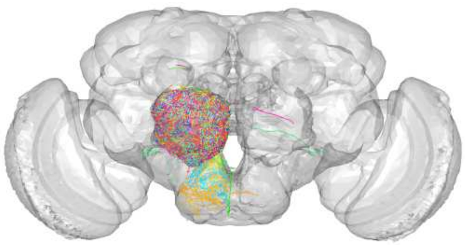

38

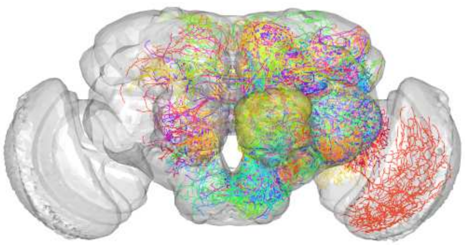

39

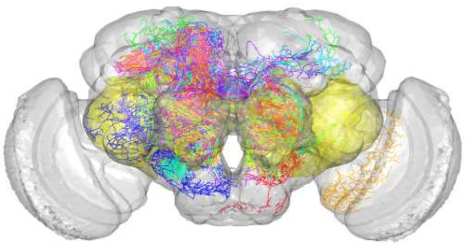

40

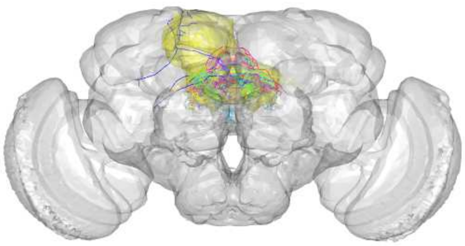

41

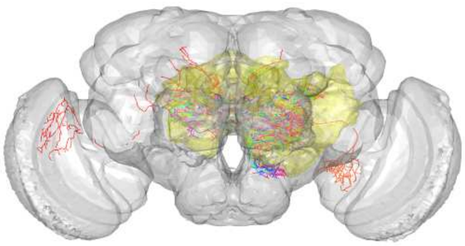

42

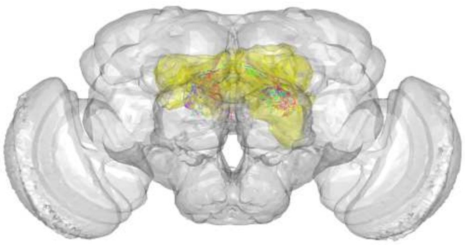

43

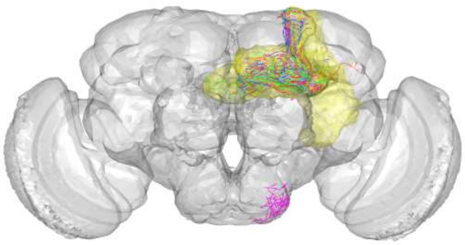

44

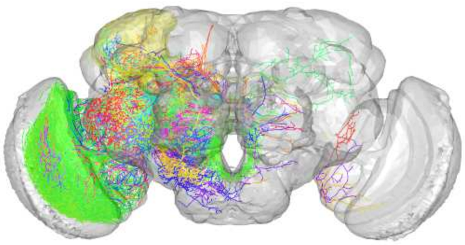

45

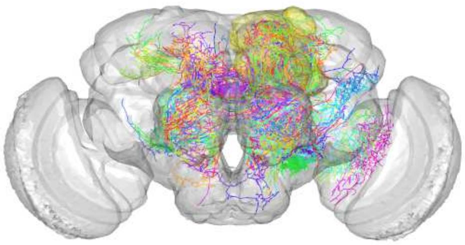

46

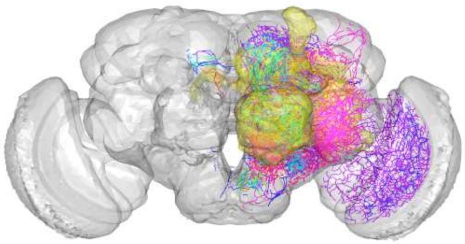

47

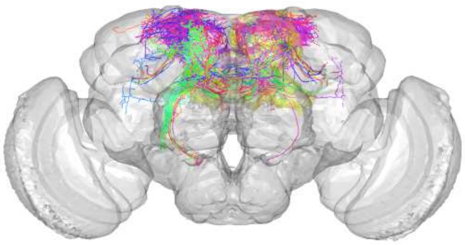

48

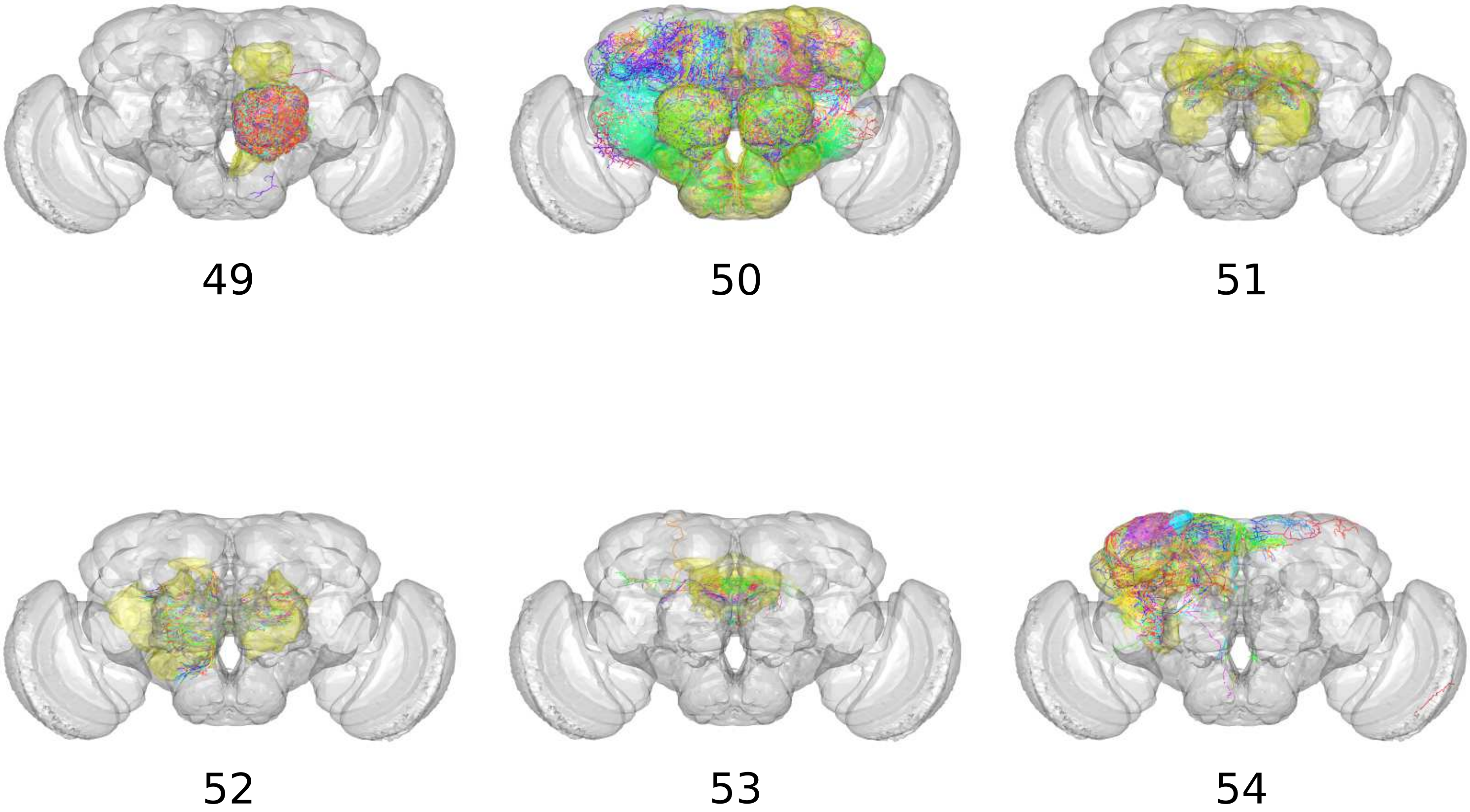


**Figure S2:** Each connectivity-based class embedded using the `natverse` 3D template of the Drosophila brain, showing its constituent neurons and innervating neuropils.
