## Supplementary material for "Circuit Analysis of the Drosophila Brain using Connectivity-based Neuronal Classification Reveals Organization of Key Communication Pathways": Figure S3.pdf

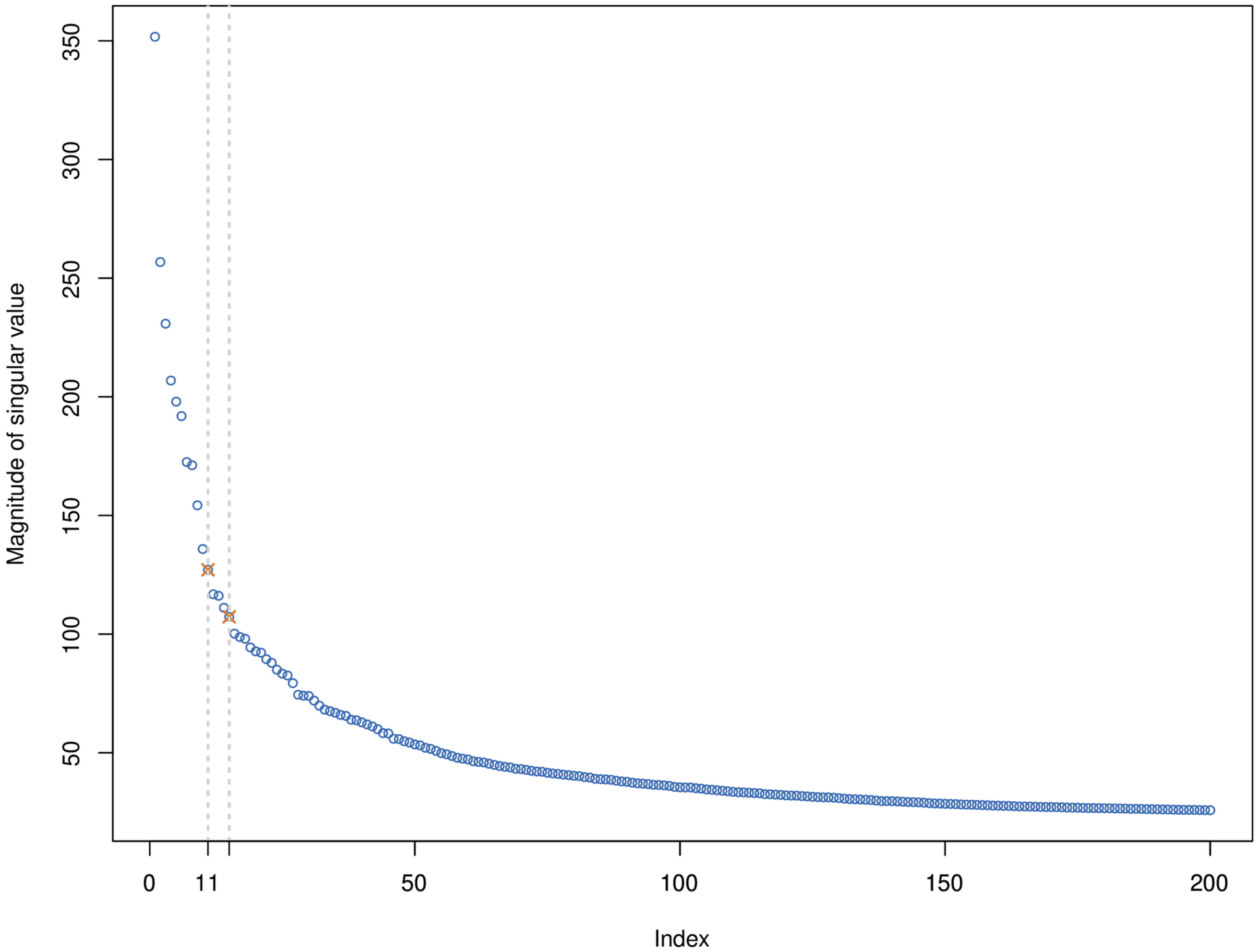


**Figure S3:** The possible choices for the embedding dimensionality *d={11,15}* were determined by identifying the first and second elbow-point (Zhu and Ghodsi, 2006), respectively, on the scree plot of singular values.
