## Supplementary material for "Circuit Analysis of the Drosophila Brain using Connectivity-based Neuronal Classification Reveals Organization of Key Communication Pathways": Figure S4.pdf

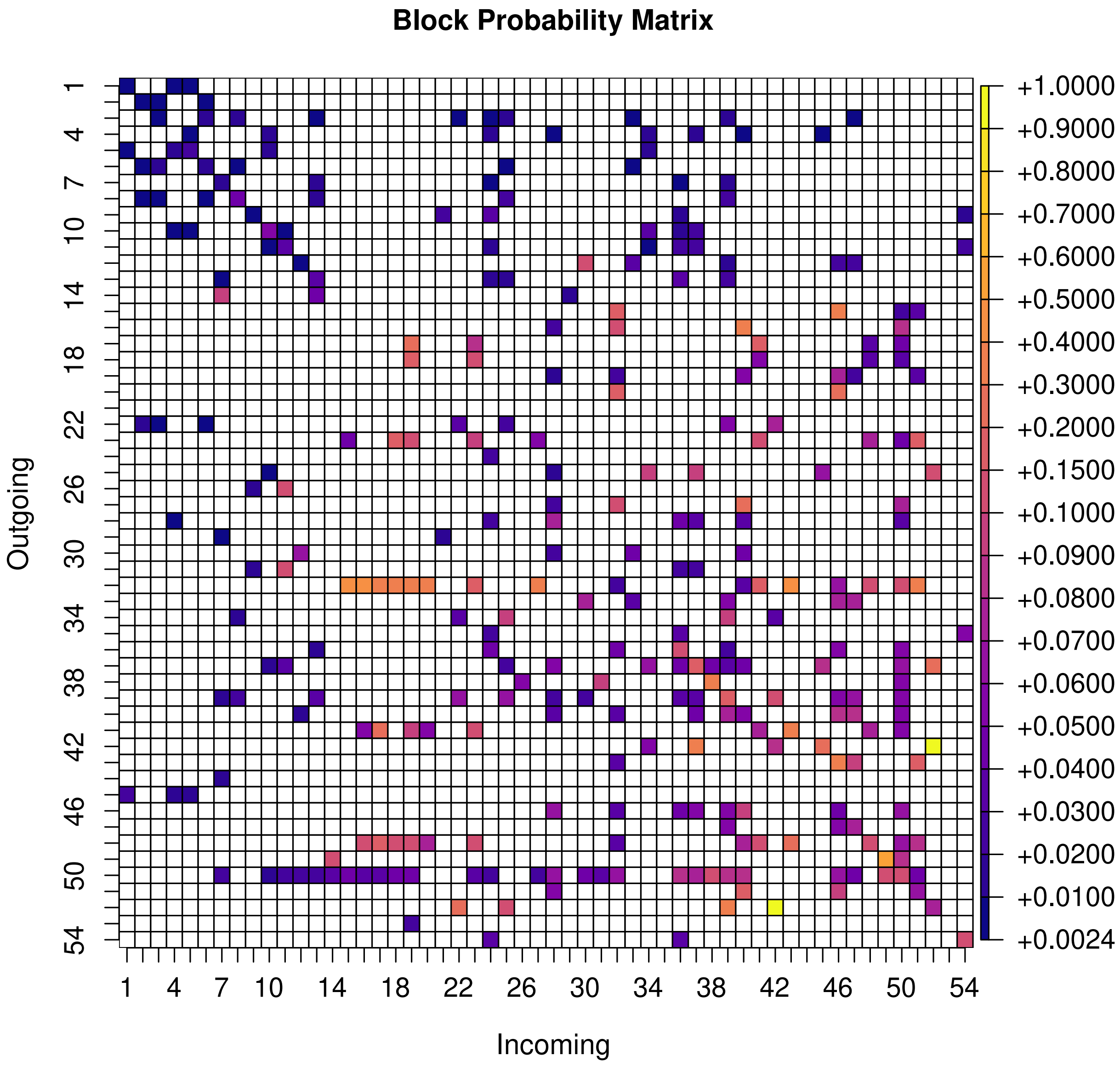


**Figure S4:** **(a)** The estimated block connectivity probabilities $\hat{p}_{ij}$'s between the 54 connectivity-based classes. The 319 non-zero $\hat{p}_{ij}$'s represent a weighted, directed edge between the 54 classes in the inferred mesoscale circuit.

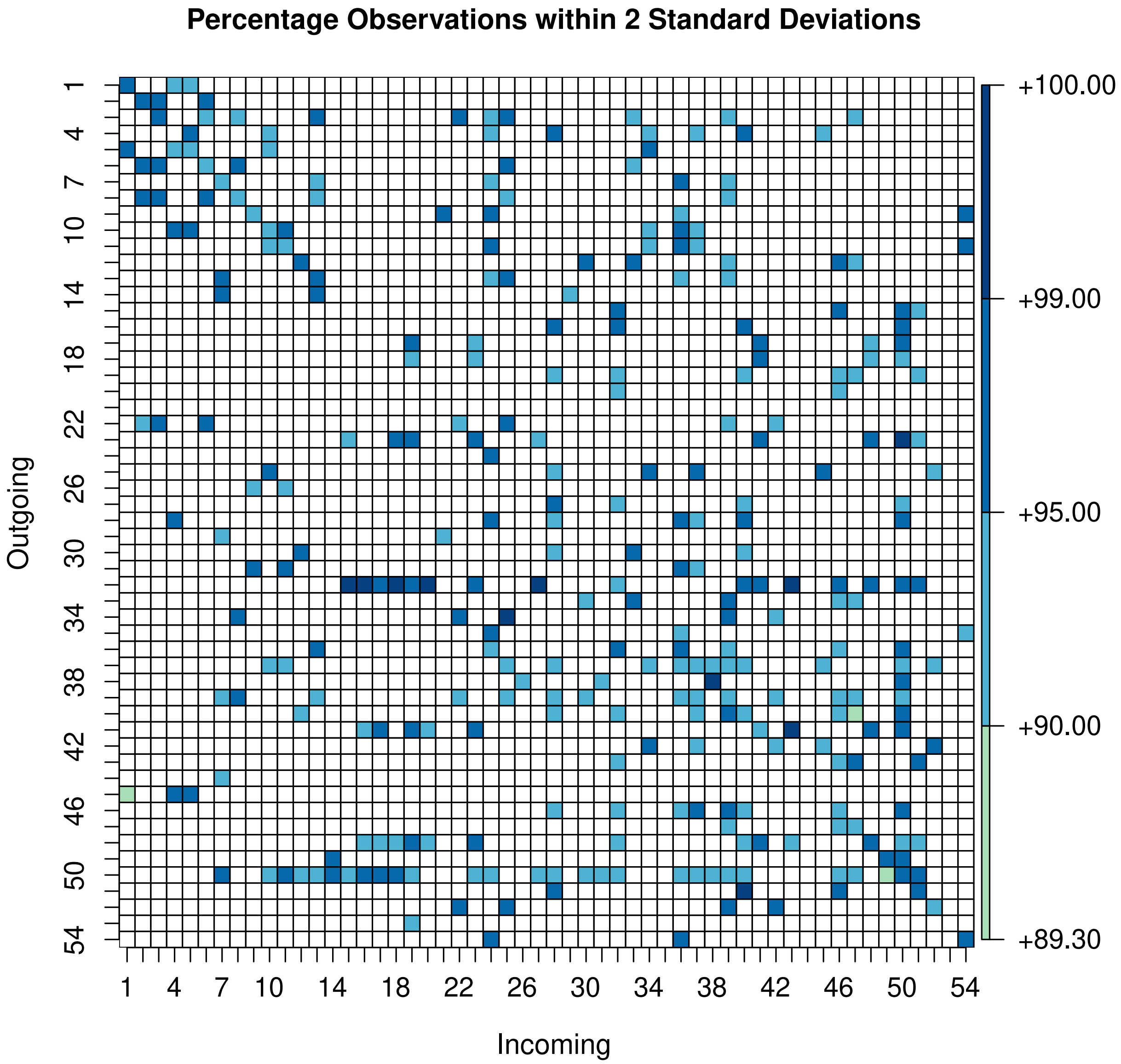


**Figure S4: (b)** The percentage of observed values for a block-pair that lie within two standard deviations of the expected binomial mean $|V_j|\hat{p}_{ij}$.

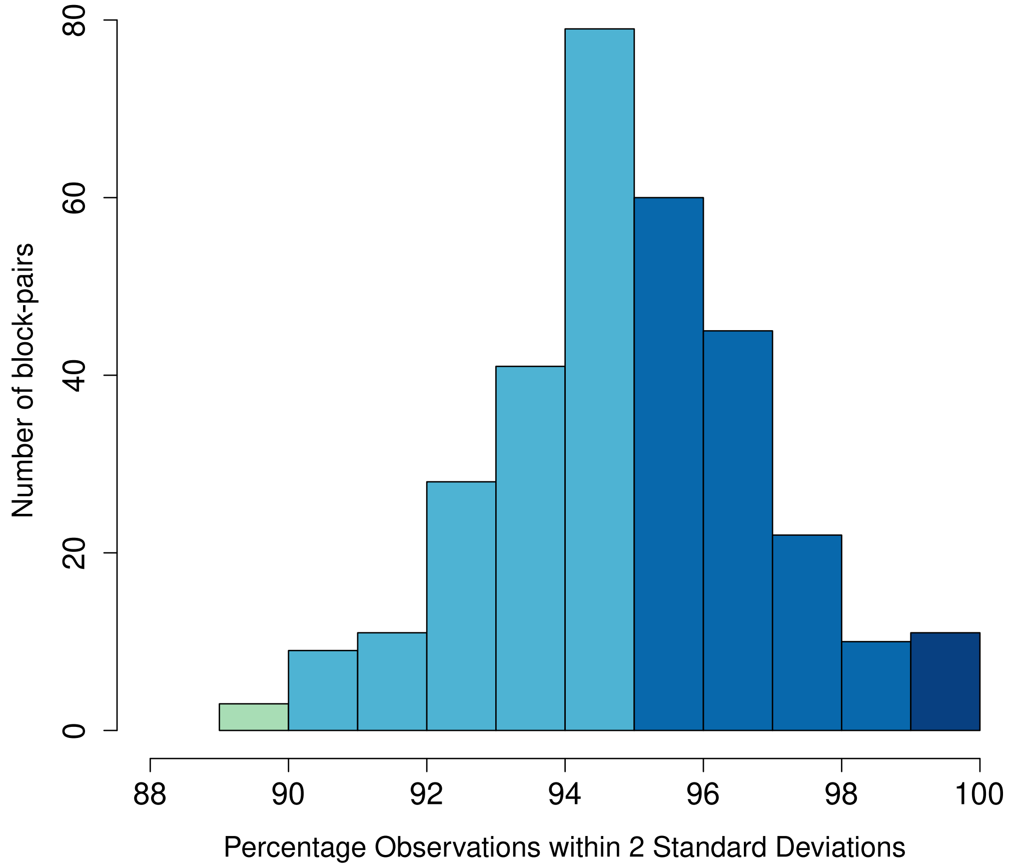


**Figure S4: (c)** A histogram showing the frequency count of block-pairs by the percentage of observed values that lie within two standard deviations of the expected binomial mean $|V_j|\hat{p}_{ij}$.
