## Supplementary material for "Circuit Analysis of the Drosophila Brain using Connectivity-based Neuronal Classification Reveals Organization of Key Communication Pathways": Figure S5.pdf

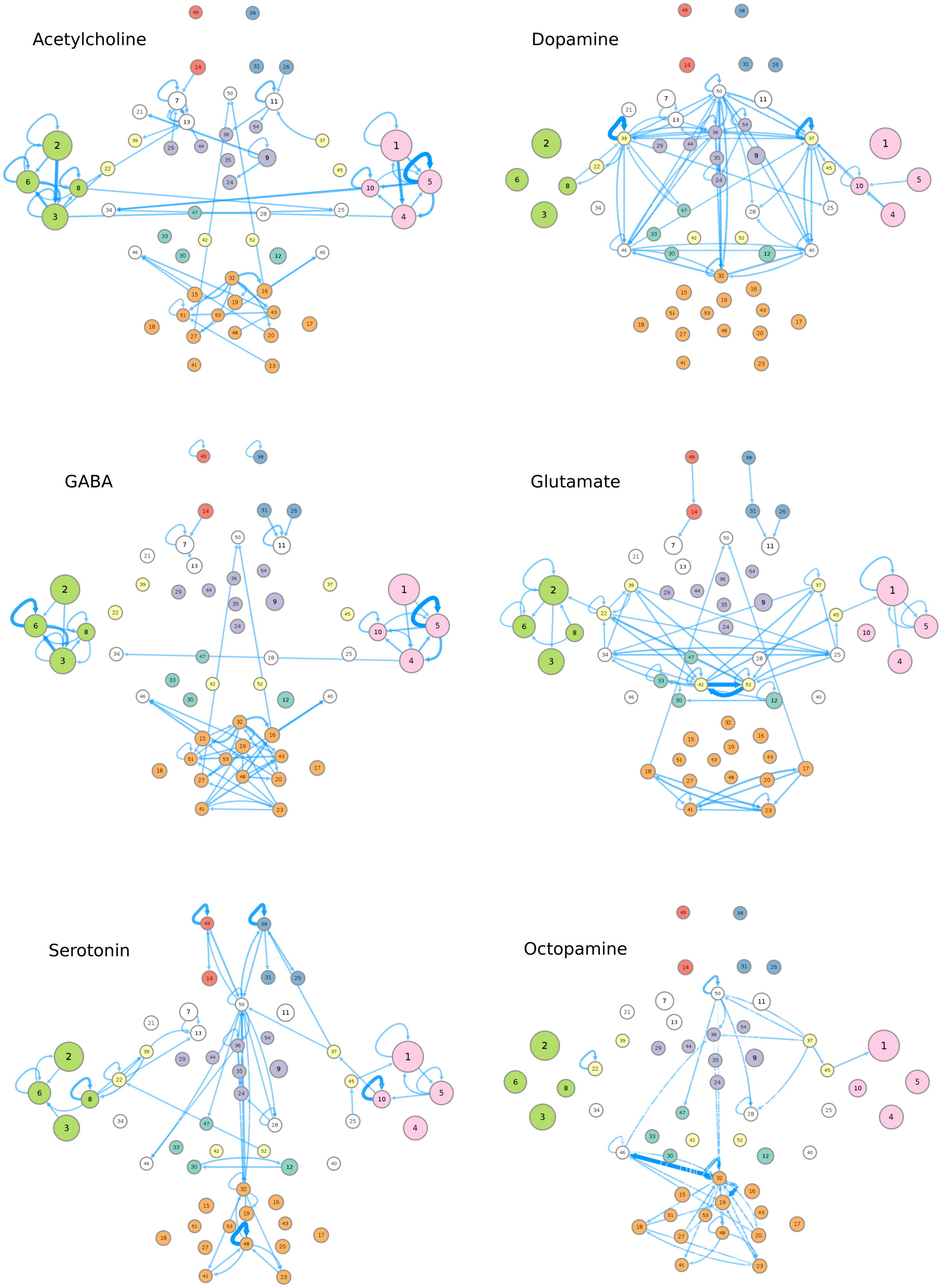


**Figure S5.** Each circuit corresponds to one neurotransmitter (NT). More specifically, to create each circuit we consider, for each class, only those neurons which are identified with that particular NT. If a particular class has zero neurons of that corresponding NT, then that class is inactive in the resulting circuit (with no incoming or outgoing edges). The edge weights are recalculated appropriately using eq. (4).
