## Supplementary material for "Circuit Analysis of the Drosophila Brain using Connectivity-based Neuronal Classification Reveals Organization of Key Communication Pathways": INFO.md

## Supplementary Figures

`Figure S1.pdf` **Random walk path length.**

`Figure S2.pdf` **3D spatial embedding of each class.**

`Figure S3.pdf` **Scree plot.**

`Figure S4.pdf` **Block connectivity probability matrix.**

`Figure S5.pdf` **Neurotransmitter pathways.**


## Supplementary Table

`Table T1.xlsx` Characterizing connectivity patterns on the circuit based on absorption and driftiness values of classes.


## Excel Files

`File F1.xlsx` **Class labels and data.**
  - Sheet *Labels*: Assigned descriptor labels for each connectivity-based class.
  - Sheet *Label Description*: Brief summary of the procedure used to determine the labels (listed in the previous sheet).
  - Sheet *data_conditional_prob*: Posterior probability $p(Y=j|X=i)$, e.g., that the neuron is associated with the j-th functional-anatomical community, given that it belongs to the i-th connectivity-based class. Also, lists the prior probabilites for comparison.
  - Sheet *data_graph_theroetic_measures*: Multiple graph theoretic measures calculated for each connecitiy-based class. Also, lists the pairwise Absorption and Driftiness values claculated using random walks on the mesoscopic circuit.
  - Sheet *data_spad*: Statistics for the neuropil (spatial) distribution of each class.
  - Sheet *data_morph+pvec*: Statistics for the morphological and persistence vector of each class.
  - The red cells highlight the column minimum. While, the green cells highlight the column maximum.

`File F2.xlsx` **ARI results.** Pairwise ARI values for different combinations of parameter choices.


## Movies

`Movie M1.mp4` **Critical growth per day.** The clip consists of 10 circuits, each corresponding to one day. This clip tracks the growth (and activity) of the circuit over the span of 10 days. If a node in the circuit has no birthed neurons until that particular day, then that node is "grayed out" and considered to be as yet inactive (unborn). For each day, the nodes that exhibit a critical growth period are colored, and so are the critical edges.

`Movie M2.mp4` **Critical growth time periods.** Similar to the S1_Movie but instead of each day, it shows circuit growth grouped into four time periods, (i) embryo, (ii) early (day1-day3), (iii) mid (day4-day6), and (iv) late (day7-day9).
